# Constricted pupil states index neural replay during wakeful rest in humans

**DOI:** 10.64898/2026.10.21.760673

**Authors:** Debadatta Dash, Leonardo Claudino, Hisato Sugata, Fumiaki Iwane, Leonardo G. Cohen, Ethan R. Buch

**Author notes:** Correspondence: Debadatta Dash, Ph.D., Tarleton State University;. Ethan R Buch, Ph.D., National Institutes of Health.

## Abstract

Neural replay has been linked to skill consolidation during wakeful rest, yet its physiological context in humans remains poorly understood. In animals, hippocampal ripples and associated replay occur more frequently within intrinsically oriented brain states associated with constricted pupils, suggesting that pupil dynamics may index neural replay events in humans. We combined magnetoencephalography-derived sequence replay with pupillometry in 25 participants learning a motor sequence across alternating intervals of practice and rest. Replay was indexed by smaller-pupil states, and replay timing clustered on the constricting phase of slow pupil fluctuations. More frequent replay accompanied larger cumulative offline gains during early learning, while smaller rest pupil was associated with better performance when practice resumed. Multivariate pupil features predicted replay-associated states across and within individuals while preceding pupil dynamics showed modest predictive ability to identify the precise onset of individual replay events. Here, we show that constricted-pupil states index neural replay during wakeful rest in humans, linking replay-associated physiology with early motor learning.

## Introduction

Neural replay, the rest-based re-expression of neural activity patterns representing previously experienced or novel action sequences, has been implicated in memory consolidation and linked with ongoing fluctuations in physiological state. In rodents, hippocampal ensembles recapitulate maze-running trajectories during awake immobility, while sharp-wave ripples organize coordinated population activity and support memory-guided behavior [1, 2]. These neural replay events are embedded within brain-wide state fluctuations [3]. During quiet wakefulness in mice, smaller pupils mark relatively low-arousal conditions with prominent slow cortical activity and higher hippocampal ripple rates [4]. During NREM sleep, recent-memory replay preferentially occurs in constricted-pupil substates, whereas older memories are replayed preferentially in dilated-pupil substates [5]. Pupil fluctuations therefore offer a noninvasive window onto physiological conditions relevant to memory processing. They covary with cortical activity and sensory responsiveness, as well as noradrenergic and cholinergic activity, while reflecting multiple autonomic influences [6, 7, 8, 9, 10]. These observations suggest that pupil-linked physiological fluctuations may help organize the expression of human memory-related neural activity.

Brief rest periods during early motor-skill learning provide an opportunity to examine this relationship in humans. In challenging motor-sequence learning, performance improvements are expressed predominantly across brief inter-practice rests, making these micro-offline gains a substantial component of early skill acquisition [11]. Understanding the neural events during these intervals is therefore central to explaining how successive bouts of practice produce skilled performance. Human neuroimaging has revealed rapid sequential reinstatement of learned representations beyond spatial navigation [12, 13], and intracranial recordings show that cortical spiking sequences recur during successful retrieval and couple to medial temporal ripples [14]. In motor learning, magnetoencephalography (MEG) has revealed temporally compressed replay of practiced sequences during short inter-practice rests, with early replay rates related to cumulative micro-offline gains [15]. Lower rest-related beta-band activity in a predominantly contralateral frontoparietal network also accompanies larger micro-offline gains [11]. These findings identify brief rest as a period of active neural processing during rapid skill acquisition, in which replay, physiological state and behavioral improvement can be examined together.

The animal pupil–ripple and sleep-replay findings suggest that physiological state may help organize memory-related activity during human wakeful rest. Emerging human intracranial evidence supports this possibility: a preprint reports higher hippocampal ripple rates during smaller-pupil wake states [16]. Yet the physiological state accompanying sequence-resolved motor replay during learning remains incompletely understood. It is unclear whether pupil-linked fluctuations observed in other replay contexts also accompany MEG-derived replay across successive practice–rest cycles, or how this state relates to learning and performance when practice resumes. A further question is whether pupil dynamics identify a replay-associated state or also carry information about the precise initiation of individual replay events.

Here, we hypothesized that neural replay during early motor-sequence learning in humans would preferentially occur during relatively smaller-pupil states within brief inter-practice rest periods. We combined MEG-derived skill sequence replay detection with simultaneous pupillometry in participants learning a motor sequence to examine pupil size and slow pupil fluctuations during replay. We assessed how replay rate and pupil state relate to early motor learning and subsequent performance, and whether multivariate pupil features distinguish replay-associated states across and within individuals and carry information about individual onset times.

## Results

### Behavioral gains, pupil dynamics and replay during learning

Correct sequence typing speed improved rapidly during early learning, with performance gains expressed predominantly across the brief inter-practice rest periods. Early learning comprised trials 1–11, by which the group performance curve approached approximately 95% of training-session improvement [17] (Fig. 1A). Mean transition-paired micro-offline gain was +0.0722 sequences/s (*p* < 0.001), exceeding micro-online gain (paired *p* = 0.029; Fig. 1C). After the group performance plateaued, negative online changes and positive offline gains largely offset each other.

**Figure 1.**
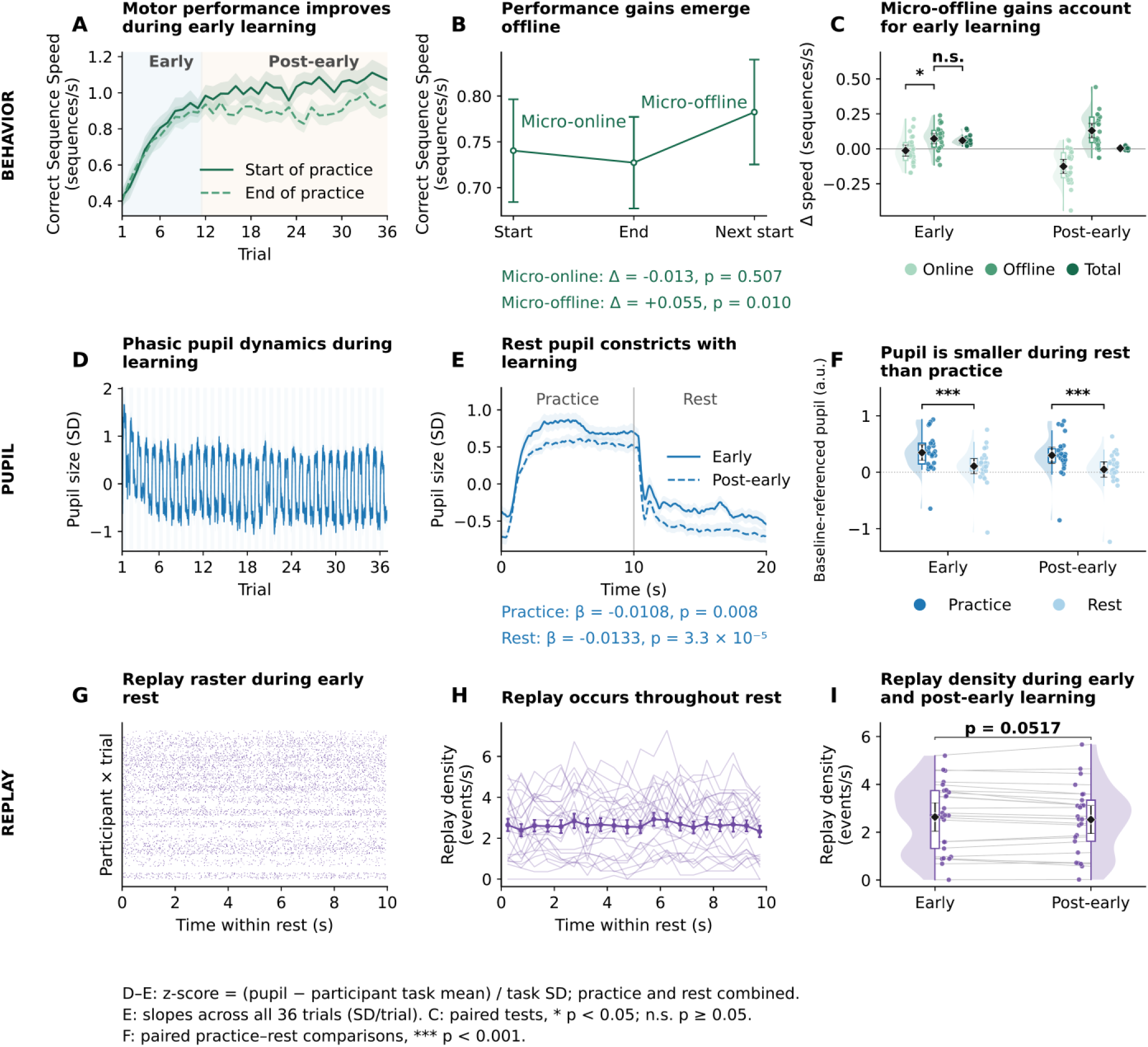
Behavior, pupil and replay during motor-sequence learning. (A) Mean correct-sequence speed at practice start and end across 36 trials. (B) Marginal start, end and next-start performance across the ten transitions linking trials 1–11 shows improvement primarily between practice bouts. (C) Transition-paired early micro-offline gain was positive (+0.0722 sequences/s, *p* < 0.001) and exceeded micro-online gain (paired *p* = 0.029); post-early online losses and offline gains largely offset one another. (D) Within-participant standardized pupil traces; shading marks practice. (E) Early and post-early practice–rest trajectories share one standardized scale; mean central-rest pupil (2–8 s) declined across training. (F) Pupil is smaller during rest than practice. Baseline-referenced trial medians gave rest-minus-practice differences of −0.240 and −0.252 a.u. in early and post-early learning (both *p* < 0.001). (G) Target-replay onsets across participants and early trials. (H) Exposure-normalized replay density across the 10-s rest interval. (I) Replay density in early and post-early learning (paired *p* = 0.052). N = 25; early, trials 1–11; post-early, trials 12–36. D/E use pooled practice/rest standardization; E trends use central 2–8-s means. In C/F/I, dots denote participants, diamonds and bars means and 95% t CIs, boxes medians and interquartile ranges, whiskers observations within 1.5 interquartile ranges, and shaded distributions densities. Bands and bars in A/B/D/E/H denote SEM. Purple denotes replay and blue pupil. CI, confidence interval; SD, standard deviation; SEM, standard error of the mean; sequences/s, sequences per second. \**p* < 0.05; \*\*\**p* < 0.001; n.s., *p* ≥ 0.05.

Pupil size varied with the practice–rest cycle and declined across training during both practice (−0.01077 SD/trial, *p* = 0.008) and rest (−0.01329 SD/trial, *p* < 0.001) (Fig. 1D–F). On the jointly filtered, baseline-referenced scale, rest pupil was smaller than practice pupil during early learning (difference −0.240 a.u., 95% confidence interval [CI] [−0.326, −0.154], *p* < 0.001) and post-early learning (−0.252, 95% CI [−0.338, −0.165], *p* < 0.001).

Replay of the practiced sequence (4–1–3–2–4) occurred across inter-practice rest periods, at a mean rate of 2.634 events/s during early learning (Fig. 1G–I; Supplementary Fig. 1). Thus, the brief rest intervals during which early behavioral gains emerged were accompanied by pronounced pupil-state transitions and sustained neural replay.

### Constricted pupil states index human motor replay

During early learning, target replay preferentially occurred when pupil size was relatively small. To reduce the influence of practice–rest visual display transition effects on pupil diameter, we focused on the central 2–8 s of each rest interval. To account for systematic pupil changes across the rest interval, we subtracted each participant’s mean pupil value at each 500-ms rest-time position. This exact time residualization tested whether pupil was smaller than usual for that participant at the same moment within rest when replay occurred. Replay-positive bins had lower residual pupil values than replay-negative bins (Δ = −0.129 SD, 95% CI [−0.249, −0.009], d_z_ = −0.445, *p* = 0.036; Fig. 2A). Central-rest (2–8 s) contrasts were −0.129 SD early, +0.004 SD post-early and −0.021 SD across all trials (Fig. 2B). The association was therefore most apparent during early learning, although the direct paired early–post-early difference remained inconclusive (−0.133 SD, 95% CI [−0.280, 0.014], *p* = 0.073).

**Figure 2.**
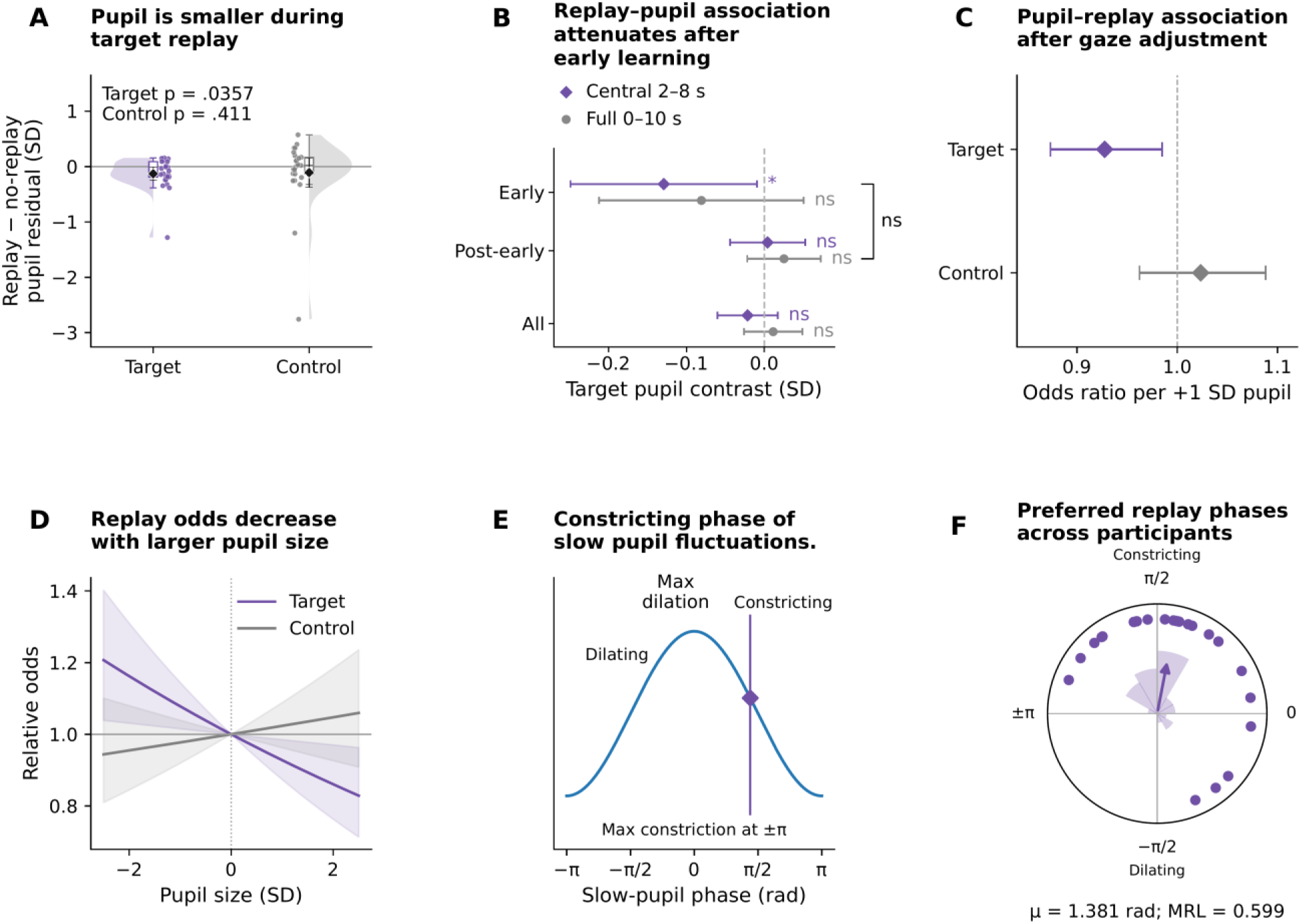
Constricted pupil states index human motor replay. (A) Exact-time-residualized pupil was smaller in target replay-positive than replay-negative bins (Δ = −0.129 SD, 95% CI [−0.249, −0.009], *p* = 0.036). Control and direct target–control comparisons are shown in Supplementary Fig. 3B. (B) Target contrasts at 500 ms for early (trials 1–11), post-early (12–36) and all trials (1–36), in central 2–8-s rest (purple diamonds) and full 0–10-s rest (gray circles). All trials pool the two periods. In B, *p < 0.05 and ns, p ≥ 0.05 denote individual two-sided contrasts against zero. The bracket spans Early and Post-early; both direct comparisons are ns (central rest p = .073; full rest p = .186). (C) Gaze- and time-adjusted target replay odds decreased with pupil size (OR = 0.928 per SD, 95% CI 0.874–0.985, *p* = 0.014). (D) Relative odds from the same model, referenced to standardized pupil = 0. (E) Observed target mean phase on the constricting phase of the schematic 0.1–1-Hz pupil cycle. (F) Participant preferred phases, rose histogram and mean resultant vector (μ = 1.381 rad, participant-bootstrap 95% circular CI 0.910–1.728 rad; MRL = 0.599; Rayleigh *p* < 0.001). Time-matched phase controls are in Supplementary Fig. 4. N = 25. A/C/D use early central 2–8-s rest; B compares early, post-early and all trials over central and full rest; E/F use full 0–10-s rest during trials 1–11. In A, points denote participants, diamonds and bars means and 95% t CIs, half violins distributions, boxes medians and interquartile ranges, and whiskers 1.5 interquartile ranges. B shows participant 95% t CIs. C uses participant-clustered normal-Wald 95% CIs; D propagates the same coefficient uncertainty. Blue denotes pupil, purple target and gray control replay. OR, odds ratio; CI, confidence interval; SD, standard deviation; MRL, mean resultant length.

Smaller pupil was associated with higher replay odds and this association survived after adjustment for linear rest time and gaze (odds ratio [OR] 0.928 per SD, participant-clustered 95% CI 0.874–0.985, *p* = 0.014; Fig. 2C,D). The association persisted in a model accounting for participant intercepts, categorical rest time, learning-trial number and gaze (OR 0.885, 95% CI 0.806–0.972, *p* = 0.011; Supplementary Fig. 3A). The control replay state contrast was not significant (−0.108 SD, p = 0.411), and the pupil contrast did not differ significantly between target and control replay (*p* = 0.883; Supplementary Fig. 3B).

The mean target contrast remained negative at 250-, 500-, 1,000- and 2,000-ms bin widths (−0.031, −0.129, −0.099 and −0.125 SD), but eligible N declined from 25 to 19 and 9 at the two coarsest widths, respectively, and uncertainty widened (Supplementary Fig. 3E). The studentized within-trial temporal-rotation test yielded *p* = 0.015 in the eligible cohort (N = 24; one participant contributed only one detected target-replay event across early rest and was not eligible; Supplementary Fig. 3C,D). Low-motion gaze restrictions reduced eligibility and precision, with no gaze-restriction comparison surviving multiplicity correction (Supplementary Fig. 5). Together with the significant exact-time and gaze-adjusted occurrence analyses, these findings support a constricted-pupil physiological state associated with neural replay.

Replay timing was also organized with respect to slow pupil dynamics. Across the full early-rest interval, participants’ preferred target-replay phases clustered during the constricting phase of 0.1–1-Hz pupil fluctuations (mean direction μ = 1.381 rad, participant-bootstrap 95% circular CI 0.910–1.728 rad, mean resultant length [MRL] = 0.599, Rayleigh *p* < 0.001; Fig. 2E,F). Control phases also showed nominal concentration. Comparisons with the same rest time in other trials did not establish replay-specific phase displacement beyond rest-time structure (Supplementary Fig. 4).

### Replay rate and pupil state track learning

More frequent and more regularly spaced replay accompanied larger offline gains during early learning. Participants with greater early target-replay rates accumulated larger offline gains during early learning (Pearson *r* = 0.484, *p* = 0.016; Fig. 3A). Lower variability in within-trial inter-event intervals, quantified by their coefficient of variation (IEI CV), was also associated with larger early offline gains (r = −0.472, *p* = 0.020; Fig. 3B). Higher replay rates accompanied more negative early online change: r = −0.481, *p* = 0.017; Fig. 3C). All three associations used the same 24 participants with estimable early replay IEI CV. The online and offline associations are related because the same end-of-practice performance measurement contributes to both changes. Rank-based sensitivities are shown in Supplementary Fig. 2B; participant-influence checks are reported in Supplementary Methods and Supplementary Data 1.

**Figure 3.**
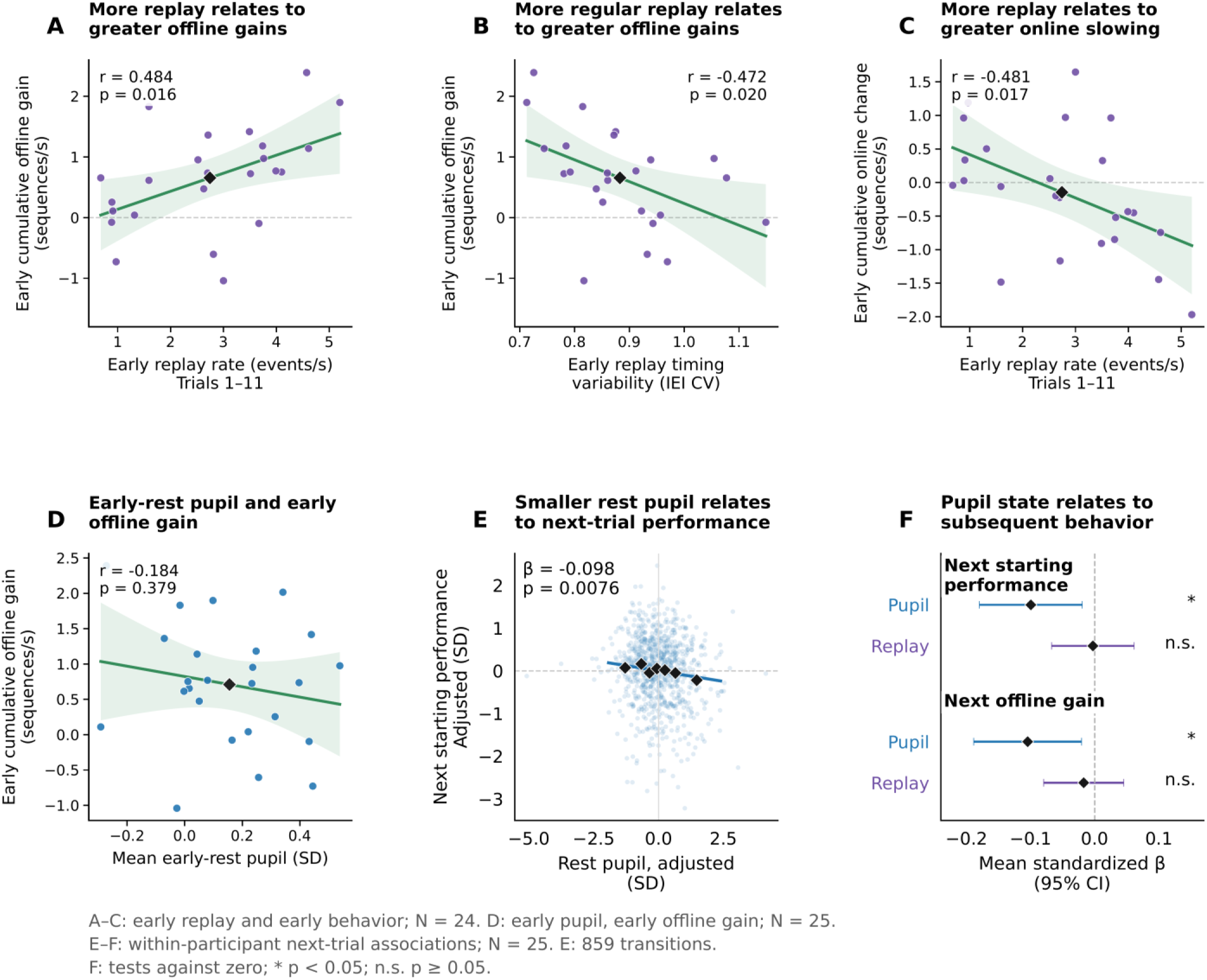
Replay rate and pupil state track learning at complementary scales. (A) Higher early replay rate accompanied larger early cumulative offline gain (Pearson r = 0.484, *p* = 0.016). (B) More regular replay relates to greater offline gains. Lower variability in within-trial replay inter-event intervals (IEI CV), indicating more regular event spacing, accompanied larger early cumulative offline gain (Pearson r = −0.472, *p* = 0.020). (C) More replay relates to greater online slowing. Higher early replay rate accompanied more negative early cumulative online change (Pearson r = −0.481, *p* = 0.017). (D) Early-rest pupil and early cumulative offline gain. Mean early-rest pupil was not significantly associated with early cumulative offline gain across participants (Pearson r = −0.184, p = 0.379). (E) Smaller rest pupil relates to next-trial performance. After adjustment for the preceding trial’s ending performance, trial, replay rate, gaze and participant fixed effects, smaller central-rest pupil was associated with better next starting performance (standardized β = −0.0983, participant-clustered *p* = 0.008; 859 transitions). The seven pupil-quantile diamonds summarize the scatter for visualization; inference uses individual transitions. (F) Pupil state relates to subsequent behavior. Participant-specific pupil coefficients were negative on average for next starting performance and next offline gain; corresponding replay coefficients did not differ significantly from zero. A–C use the common N = 24 sample with estimable early IEI CV; D uses N = 25. E/F are within-participant analyses (N = 25). Early replay and online change cover trials 1–11; early offline gain sums the ten transitions linking trials 1–11. D compares early pupil with early offline gain. In A–D, points represent participants, diamonds mark bivariate means, lines show linear fits and shading denotes 95% CIs for the mean response. D/E/F use central 2–8-s; Pupil variables were standardized within participant. Panel E includes all transitions; in F, diamonds and bars denote means and 95% t CIs, with tests against zero. Purple denotes replay, blue pupil and green behavioral fits. IEI CV is interval SD/mean; CI, confidence interval; SD, standard deviation; sequences/s, sequences per second. \**p* < 0.05; n.s., *p* ≥ 0.05.

Mean early-rest pupil was not significantly associated with early cumulative offline gain across participants (r = −0.184, p = 0.379, N = 25; Fig. 3D), indicating that participants with smaller average pupils during early rest did not show a statistically clear tendency to accumulate larger early offline gains. Within participants, however, constricted pupil during rest was associated with better performance when practice resumed. This relationship remained after accounting for the preceding trial’s ending performance, trial, replay rate, gaze and participant fixed effects (standardized β = −0.0983, participant-clustered 95% CI [−0.1705, −0.0261], *p* = 0.008; 859 transitions, 25 participants; Fig. 3E). A wild-cluster sensitivity supported the association (*p* = 0.013), and participant-specific pupil coefficients were negative on average for next starting performance and next offline gain (Fig. 3F). Replay rate thus tracked individual differences in learning, whereas pupil state tracked fluctuations in subsequent performance within individuals.

### Pupil dynamics distinguish replay-associated states

Multivariate summaries of pupil level, variability and gradient distinguished replay-associated from temporally matched inactive task samples during central rest (2–8 s). In nested cross-validation, mean participant area under the receiver operating characteristic curve (ROC AUC) was 0.857 for held-out participants in leave-one-subject-out (LOSO) evaluation and 0.816 for held-out trials within individuals (N = 24; Fig. 4A).

**Figure 4.**
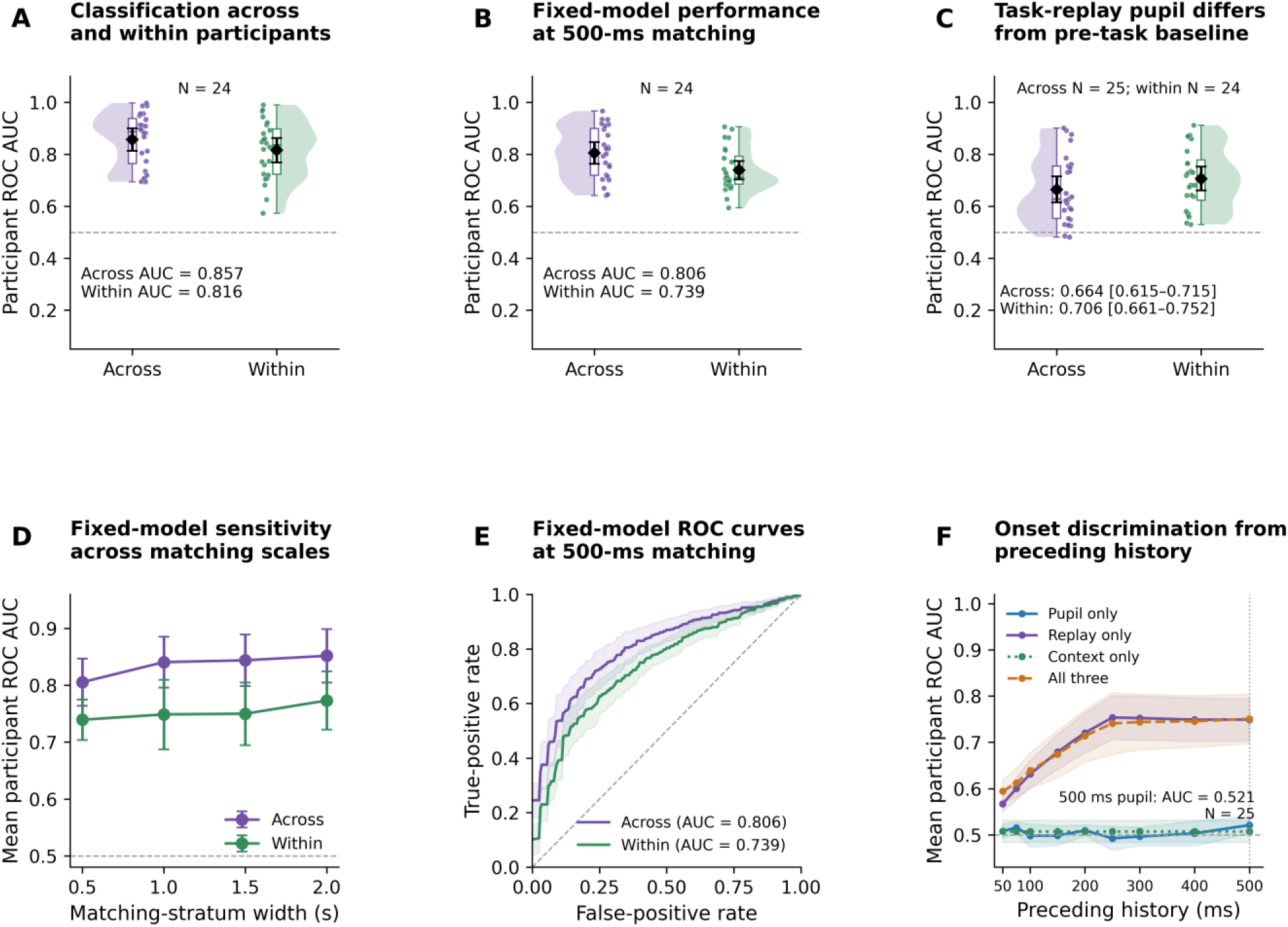
Pupil representations distinguish replay-associated states across and within participants. (A) Nested classification of replay-associated versus temporally matched inactive central-rest states across held-out participants (LOSO ROC AUC = 0.857) and within participants across held-out trials (AUC = 0.816; N = 24). (B) With linear detrending, the extended-feature representation and LDA held fixed, 500-ms matching yielded AUC = 0.806 across participants and 0.739 within participants (N = 24). (C) Replay-associated task pupil versus pre-task pupil (across AUC = 0.664, N = 25; within AUC = 0.706, N = 24), averaged over 199 baseline allocations. Pre-task rest preceded sequence experience and provides a physiological reference; behavioral context and recording conditions differed. (D) Fixed-model sensitivity across 0.5–2.0-s matching strata (N = 24). (E) Participant-average ROC curves at 500-ms matching (N = 24). A–E select samples from 2–8 s after applying the original whole-rest transforms. (F) Histories preceding candidate replay/control samples carried modest pupil-only onset information at 500 ms (AUC = 0.521, 95% CI 0.503–0.540; nine-window BH q = 0.0622) and stronger replay-history information (AUC = 0.749; N = 25). A/B/D/E classify sampled task states; C compares task and pre-task pupil; F classifies onset times. These are different prediction tasks. Dots in A–C denote participants, diamonds means, half violins distributions, boxes medians and interquartile ranges, and whiskers observations within 1.5 interquartile ranges. A/B/D–F show participant 95% t intervals, pointwise in E/F; C shows conditional participant-bootstrap 95% intervals. Dashed lines denote chance. ROC AUC, area under the receiver operating characteristic curve; LOSO, leave-one-subject-out; BH, Benjamini–Hochberg; CI, confidence interval.

We next examined how discrimination varied with the temporal resolution used to match replay-associated and inactive samples within rest. With the linear discriminant analysis (LDA) pipeline held fixed, 500-ms matching yielded AUCs of 0.806 across participants and 0.739 within participants (Fig. 4B,E). Performance increased as matching strata became broader (Fig. 4D). Model-selection frequencies are documented in Supplementary Fig. 8.

Pupil features from replay-associated central-rest periods also distinguished task rest from pre-task rest (Fig. 4C). Nested across-participant AUC was 0.664 (conditional participant-bootstrap 95% interval 0.615–0.715; N = 25), and nested within-participant AUC was 0.706 (0.661–0.752; N = 24). Pre-task rest provided a physiological reference before sequence training, although differences in behavioral and recording context prevent a replay-specific interpretation (Supplementary Fig. 6, Supplementary Table 1 and Supplementary Data 6). Full-rest and central-rest results, including a sensitivity that recomputed transforms within central rest, are compared in Supplementary Table 4.

We next asked whether preceding pupil measurements contained information about the start of individual replay events. At the designated 500-ms history window, ending immediately before each candidate replay or control sample, pupil history carried modest event-start information (AUC = 0.521, 95% CI 0.503–0.540, one-sided *p* = 0.014, nine-window BH q = 0.0622; N = 25). Recent replay history carried stronger timing information (AUC = 0.749; Fig. 4F). The pupil-only effect did not survive correction across the nine tested history windows (50–500 ms; Supplementary Fig. 7 and Supplementary Table 2). State and onset classification address different sampling and prediction tasks; their AUCs are therefore not directly comparable measures of the same discrimination problem. In sum, pupil features characterized replay-associated states but carried little information about exact onset timing.

## Discussion

Human motor replay occurred within a measurable pupil-linked physiological context during wakeful rest. Replay was enriched during smaller pupil states, and its timing clustered on the constricting phase of slow pupil fluctuations. Pupil therefore provides an accessible physiological correlate of sequence-resolved replay while participants learn a motor skill. The association spans pupil level, occurrence models and multivariate state discrimination, connecting brief replay events to pupillary fluctuations in the state of the learner. Human motor replay thus occurred in a pupil-linked state that also tracked early motor learning.

### A pupil-linked physiological context for replay

Previous work links pupil state to hippocampal ripples during quiet wakefulness and to the organization of replay during sleep. In awake mice, small pupils accompany hippocampal ripples [4]; during NREM sleep, pupil substates organize replay of recent and older memories [5]. Emerging human evidence complements these observations: an intracranial preprint reports more hippocampal ripples during small-pupil wake states [16], and cortical sequence reinstatement can couple to medial temporal ripples during retrieval [14]. The present findings extend this physiological perspective to MEG-derived motor-sequence replay during brief inter-practice rest. Pupil-linked cortical fluctuations also accompany changes in sensory responsiveness and neuromodulatory activity [6, 7, 8, 10]. Noradrenergic and cholinergic activity are therefore potential influences on the state accompanying replay [7, 8], although pupil measurements alone cannot identify a single pathway. Across these settings, pupillometry provides a common measure for examining how memory-related neural activity varies with physiological state.

A slowly varying physiological state and a brief replay event need not share the same time course. A pupil-linked state may span multiple events without changing at each onset. Across the full early-rest interval, participants’ preferred replay phases clustered during pupil constriction, relating event timing to an ongoing slow fluctuation. The classification analyses address the distinction between state and event timing: multivariate summaries distinguished replay-associated samples, whereas preceding pupil measurements carried modest information about individual event starts. Increased state discrimination with broader matching strata is compatible with broader temporal structure contributing to the sampled-state comparison. Because state and onset classification use different sampling and prediction tasks, however, their AUCs do not directly measure the same temporal information. Task–baseline discrimination provides a further physiological reference, showing that replay-associated task samples differ from pupil measurements before sequence training.

### Replay and pupil state link waking rest to motor learning

Replay rate and pupil state relate to motor learning at complementary levels. The pupil–replay association was most apparent during early learning, when substantial micro-offline gains were expressed, suggesting a potential relationship between replay-associated physiological state and early skill acquisition. Participants with more frequent and less variably spaced early replay accumulated larger early offline gains. Greater replay rates also accompanied more negative early online change. The online and offline associations are related because the same end-of-practice measurement contributes to both; they describe components of the same early learning process. Mean early-rest pupil, by contrast, showed no clear relationship with cumulative early offline gain across participants. Within participants, smaller rest pupils accompanied better performance when practice resumed, after adjustment for preceding performance and other covariates. These observations distinguish individual differences in accumulated improvement from fluctuations in subsequent performance across successive practice bouts.

The within-participant pupil–performance relationship may reflect changes in arousal or task engagement as practice resumes. Other physiological measurements also connect brief rests to early learning: lower 16–22-Hz beta activity in a predominantly contralateral frontoparietal network accompanied larger micro-offline gains [11]. Replay, pupil state and oscillatory activity thus offer complementary perspectives on these intervals. The behavioral distinction matters: a physiological measure that tracks performance from one practice bout to the next need not also explain why some participants accumulate larger gains than others.

### Limitations

Across bin-level comparisons, gaze-adjusted occurrence models, slow-phase analyses and multivariate classification, replay accompanied a smaller pupil physiological state that also tracked subsequent performance within individuals. These converging associations link replay, pupil state and learning without identifying a causal mechanism. Target replay was significantly associated with pupil state and behavior, and control replay was not. However, the direct target–control contrast in pupil state did not establish a sequence-specific difference. Pupil size also declined across training, potentially reflecting fatigue, engagement, attention or habituation. Focusing the replay–behavior analyses on early learning reduced exposure to accumulated time-on-task effects. Exact-time residualization accounted for the average within-rest pupil profile, and adjustment for learning-trial number addressed systematic changes across training. These steps may partly account for fatigue-related changes that covary with time on task, but fatigue and vigilance remain unresolved without independent measurements. Bidirectional zero-phase filtering also means that preceding-history classification supports retrospective temporal discrimination rather than online forecasting. Independent replication will be important for establishing how broadly these relationships generalize. Pupil size reflects broad autonomic and neuromodulatory influences. Establishing a causal link between a specific physiological state and neural replay is therefore difficult with pupillometry alone.

Pupillometry offers a noninvasive window onto the physiological context accompanying human motor replay. By linking neural activity during brief waking rests with the state of the learner, it provides a practical approach to studying replay as motor skills develop.

## Methods

### Participants

Thirty-three participants were enrolled in the parent study. Two were excluded because of technical problems during MEG acquisition, leaving 31 participants in the released dataset [17, 18]. For the present multimodal pupil–replay analysis, six participants were excluded because of absent or unusable replay detections, unusable pupil recordings, or pupil–task alignment problems, leaving 25 participants in the primary cohort (Supplementary Table 3). Sample size was determined by the availability of participants from the existing dataset who met the multimodal data-quality and analysis eligibility criteria; no prospective power calculation was performed for this reanalysis. Participants were healthy right-handed adults who provided written informed consent; the study was approved by the Combined Neuroscience Institutional Review Board of the National Institutes of Health [17].

### Experimental task

Participants repeatedly typed the five-element sequence 4–1–3–2–4 as quickly and accurately as possible with the non-dominant left hand using a Cedrus LS-LINE response pad (Cedrus Corp.). Keys 1–4 corresponded to the little, ring, middle and index fingers, respectively. Individual keypress identities and times were recorded throughout the experiment [17].

A six-minute pre-training rest period preceded 36 practice–rest cycles. Each cycle comprised 10 s of sequence practice followed by 10 s of inter-practice rest. During practice, the sequence remained visible and asterisks indicated the participant’s position within it without providing correctness feedback. During inter-practice rest, five X symbols replaced the sequence and participants were instructed to maintain fixation. During pre-training rest, participants fixated a central cross. The analyses included both training and pre-training rest.

Early learning was defined as trials 1–11, the interval over which approximately 95% of training-session learning was reached at the group level based on an exponential fit to the group learning curve [17]. Post-early learning comprised trials 12–36.

### MEG acquisition and replay estimation

MEG was recorded continuously at 600 Hz with a whole-head 275-channel CTF system during pre-training rest and sequence training. Replay estimates were obtained using the published sequence-decoding framework [15]. Briefly, classifiers trained on practice-related finger representations were applied to resting MEG activity to identify temporally compressed reinstatement of five-element sequences. Replay detections were represented as binary time series sampled at 200 Hz during each 10-s inter-practice rest interval. The replay detections analyzed here corresponded to the approximately 50-ms sequence duration identified in the parent replay analysis [15].

Target replay represented the practiced sequence, 4–1–3–2–4. The control sequence, 3–3–4–3–3, was the structurally unrelated control used in the parent replay analysis [15]. A replay onset was a transition from inactive to active state in the binary raster; an active first sample counted as an onset.

### Pupillometry acquisition and preprocessing

Eye position and pupil size were recorded monocularly at 600 Hz with an EyeLink 1000 Plus eye tracker (SR Research, Canada), concurrently with the behavioral task. Pupil size and horizontal and vertical gaze coordinates were retained. Pupil size was analyzed in arbitrary units (a.u.) and was not interpreted as absolute physical pupil diameter. Subsequent analyses used within-participant normalization or baseline-referenced contrasts as specified below.

Continuous pupil data were preprocessed before epoching using the open-source Kret–Sjak-Shie pupil-preprocessing pipeline [19, 20]. The pipeline rejected out-of-range samples, dilation-speed outliers, gap-edge artifacts and isolated segments, followed by iterative rejection of deviations from a smooth reconstructed trace. Median-absolute-deviation thresholds provided robust outlier criteria [21]. The final smooth reconstruction supplied the filtered pupil signal.

The filtered continuous trace was used for all subsequent pupil analyses, with normalization defined separately for each analysis. Task and pre-task recordings used for baseline comparisons were processed jointly on a common scale. The first 300 s of pre-training rest, for which valid coverage was available across participants, served as the pre-task physiological reference.

### Epochs and normalization

Practice starts were identified from task-marker transitions and used to extract 36 complete 10-s practice and 10-s rest cycles per participant. Pupil was aligned to the 200-Hz replay series by retaining every third 600-Hz sample. Trials in which both replay rasters were continuously active throughout rest were treated as invalid and excluded from replay analyses and the behavioral model; all other valid trials were retained. Primary replay–pupil state analyses focused on the central 2–8 s of each rest interval to minimize pupil dynamics associated with transitions into and out of the rest period.

Normalization was analysis-specific. Descriptive task trajectories used the participant’s pooled practice-plus-rest mean and sample SD. Because one mean and SD were applied to the entire task record within each participant, this standardization rescales the pupil trace but preserves the direction and relative shape of within-session temporal trends. Baseline-referenced contrasts used the mean of trial-level 10-s medians from the jointly filtered task trace minus the median of all finite samples in the first 300 s of filtered baseline. For the between-participant early-rest pupil analysis, pupil was standardized over all rest samples within participant (Fig. 3D). Exact-time state effects removed the across-trial pupil profile and used a bin-level sample SD. Onset-classification features used pooled practice/rest population-SD standardization.

### Behavioral measures and pupil trends

Motor performance (skill) was quantified using the correct-sequence-typing speed. Each value represents the inverse duration, in seconds, of a complete correctly performed five-key sequence and is expressed in sequences per second [17]. The first and last correct-sequence-speed samples in each practice trial defined starting and ending performance. If a practice trial contained no valid correct-sequence-speed sample, starting and ending performance were treated as missing. Micro-online gain was ending minus starting speed. Micro-offline gain was the next trial’s starting speed minus the preceding trial’s ending speed. Cumulative gains summed available changes within the specified analysis window. For Fig. 3A,B, early offline gain summed the ten transitions linking trials 1–11, excluding trial 11→12; early online change in Fig. 3C summed ending-minus-starting speed over trials 1–11. For the marginal comparison of mean micro-online and micro-offline gains during early learning (Fig. 1B), starting, ending and next-start performance were averaged separately across the ten adjacent transitions linking trials 1–11, and contrasts were then formed from those marginal means. For the transition-paired analysis of gains across learning periods (Fig. 1C), changes were calculated for each eligible transition first and then averaged within participant for the early or post-early period.

Pupil trends were calculated by regressing each participant’s central 2–8-s mean during practice or rest on trial number across all 36 trials. The participant slopes were tested against zero. Early replay rate was calculated from valid 10-s rest exposure in trials 1–11. Replay temporal regularity during early learning was quantified from within-trial inter-event intervals. Intervals were pooled across early trials containing at least three replay onsets, and IEI CV was calculated as the sample SD divided by the mean interval; participants were required to contribute at least three pooled intervals (Fig. 3B). One participant from the original cohort did not satisfy this criterion with insufficient early events and thus was excluded in relevant analyses; this analysis-specific choice did not change the N = 25 physiological or within-participant analyses.

### Within-participant behavioral models

We constructed the 35 adjacent trial transitions per participant and retained complete observations for outcome and covariates. Continuous pupil, replay, gaze, and behavioral variables were standardized within participant after complete-case selection, while trial number retained its original scale. An ordinary least-squares model related next starting speed to previous ending speed, trial, central-rest pupil, central-rest replay rate, mean horizontal and vertical gaze, median gaze-step speed, gaze dispersion, gaze path length, and participant fixed effects. Standard errors were clustered by participant with the finite-sample correction. The pupil coefficient was additionally tested with 4,999 participant-level sign flips of restricted-model residuals, using a two-sided studentized wild-cluster statistic. Participant-specific regressions used the same covariates and separately modeled standardized next starting speed and offline gain. For each complete valid rest raster, replay onsets within [2,8) s were counted and divided by six seconds to obtain the central-rest replay rate. Complete-case selection retained finite standardized outcomes and covariates; invalid replay inputs were excluded.

### Pupil state and replay occurrence

Rest pupil was averaged into twenty 500-ms bins per trial. For each participant, the mean pupil value at each absolute 500-ms bin position was calculated across all 36 trials and subtracted from that position’s trial values. The residual was divided by the participant’s sample SD across all rest-bin values; negative residuals indicate smaller pupil than typical for that participant at the same time within rest. Bins 4–15, corresponding to 2–8 s, were selected from valid early trials. A bin was replay-positive if it contained an onset. For each participant, the state effect was the mean residual pupil in replay-positive bins minus that in replay-negative bins. Target and control rasters were analyzed separately, with a paired participant test of their difference. The 500-ms scale was selected to characterize the slower pupil state surrounding brief replay activity rather than the isolated approximately 50-ms neural replay event. Sensitivity analyses repeated the comparison with 250-, 1,000- and 2,000-ms bins (Supplementary Fig. 3E). To assess sensitivity to learning period and rest window, the 500-ms target contrast was also calculated for early (1–11), post-early (12–36) and all trials (1–36) and for central 2–8-s versus full 0–10-s rest (Fig. 2B). The all-trials estimate pools both periods. Paired sensitivities compared participant effects in early trials 1–11 with post-early trials 12–36 using the same normalization, eligibility and 500-ms bins (Supplementary Methods).

Temporal controls rotated the 12 central residual-pupil bins within each valid early trial while keeping replay labels fixed, preserving trial means and replay-bin counts. The 20,000 draws included identity rotations and shared offsets across target and control rasters. For the studentized statistic, two-sided empirical *p* values were calculated from the corresponding conditional reference distributions using absolute deviations from zero and a plus-one correction (Supplementary Fig. 3C,D).

The gaze-adjusted binomial-logit model related replay occurrence to pupil, linear absolute rest time and five gaze measures, with participant-clustered uncertainty. Pupil and gaze were standardized within participant over eligible early central bins. The combined time-and-gaze sensitivity included pupil, participant intercepts, categorical within-rest bin, learning-trial number and the same five gaze covariates, replacing the linear time term with categorical bins. Both models used the same pupil/gaze construction: pupil was first referenced to the pooled practice/rest mean and sample SD, after which pupil and gaze were sample-z-scored across eligible bins within participant. Both models included 3,120 bins from 260 trials in 25 participants, using identical standardization. Participant-clustered covariance estimates used the finite-sample correction, and 95% CIs were calculated using the normal-Wald approximation. Benjamini–Hochberg (BH) correction covered the two target/control pupil tests separately for each model family; the combined model was treated as an additional occurrence sensitivity. These coefficients combine within- and across-trial variation. For visualization of the pupil effect, model-implied relative odds were calculated as exp(βx), where x denotes standardized pupil size and x = 0 is the reference; confidence intervals were propagated from fitted coefficient uncertainty (Fig. 2D).

### Slow pupil phase

Slow phase was computed from each continuous filtered pupil trace after trimming terminal nonfinite samples. A fourth-order 0.1–1-Hz Butterworth filter in second-order-section form was applied forward and backward with odd padding of 27 samples, followed by the Hilbert transform. Circular means were first calculated within participant; a Rayleigh test then evaluated the distribution of participants’ preferred phases. Primary target and control phase tests were unadjusted. Level-based replay–pupil analyses focused on central rest to minimize practice–rest transition dynamics, whereas phase analyses used the full rest interval to characterize replay with respect to the complete slow pupil cycle. Uncertainty in the group mean direction was summarized with a participant-level circular percentile bootstrap (100,000 resamples). Each resample drew 25 participant-specific preferred phases with replacement; the 2.5th and 97.5th percentiles of angular deviations from the observed circular mean defined the 95% interval. Phase concentration was quantified using the mean resultant length (MRL), which ranges from 0 for uniformly distributed phases to 1 for perfect phase alignment.

The phase-matching analysis selected the same absolute rest time from the two nearest eligible early trials other than the replay trial, ranking candidates by trial distance and breaking ties toward the earlier trial. Control windows were excluded if same-raster replay occurred within ±50 ms, and both matched controls were required to meet this criterion. Event-level complex-vector differences were averaged within participant, with at least two matched events required. Two-dimensional Hotelling tests assessed participant-level phase-vector differences separately for target and control replay in the full and central rest windows; BH correction covered these four tests.

### Pupil state classification

Samples were restricted to central rest [2,8) s after preprocessing and transformation of the complete 10-s rest trace. The absolute-time matching grid was intersected with this interval. Within each participant and trial, for temporal strata containing both classes, all target-replay-positive pupil samples were retained and an equal number of no-target-replay samples were selected from the same stratum, with replacement only when necessary. Matching-stratum width specifies the temporal region within rest from which replay-associated and inactive task samples were matched. Replay-associated and matched no-replay feature vectors from the same trial remained together in cross-validation. Each eligible trial contributed one replay-associated feature vector and one matched comparison vector. At least ten genuine trial pairs were required, retaining 24 participants and 835 trial pairs (excluding the same participant with one early replay event). Nested across-participant LOSO evaluation left out one participant, with three participant-grouped inner folds for model selection. The 36 candidates crossed 500-, 1,000- and 2,000-ms matching strata, direct trial standardization or linear detrending followed by standardization, eleven pupil features or 34 pupil-and-gradient features, and L1 logistic regression, L2 logistic regression or linear discriminant analysis (LDA). Feature scaling was fitted within training folds. The model with the highest mean inner-fold AUC was refitted to all outer-training observations.

Nested within-participant evaluation selected models separately for each participant using five outer and three inner GroupKFold splits, grouping whole trial pairs. The same candidate grid and selection rule were used. Outer predictions were concatenated to calculate participant AUCs. These validation designs evaluate transfer to unseen participants and discrimination across held-out trials within an individual, respectively.

To characterize matching-scale effects with architecture held constant, preprocessing, feature family and classifier were fixed to the original full-rest reference pipeline: linear detrending followed by trial standardization, the 34-feature pupil-and-gradient representation and LDA. This model was evaluated at 500-, 1,000-, 1,500- and 2,000-ms matching widths; 1,500 ms was evaluated only in this fixed-model sensitivity. Across-participant evaluation used LOSO; within-participant evaluation used five whole-trial-pair grouped folds. Mean ROC curves were calculated by equally weighting participant-level ROC curves. AUC ranked class scores within participant, paired accuracy measured how often the replay vector scored higher than its matched comparison, and balanced accuracy averaged class sensitivities at a probability threshold of 0.5. Exact feature definitions and estimator settings are given in Supplementary Methods; model-selection and held-out performance records are provided in Supplementary Data 5. Replay labels defined the sampled task states, so this analysis assessed retrospective state discrimination.

### Task-replay pupil versus pre-task baseline pupil

For the task–baseline comparison, pupil traces from inter-practice rest and pre-task rest were jointly processed for all 25 participants. Each eligible trial contributed one feature vector from target-replay-active task samples within [2,8) s and one measured baseline feature vector with the same relative sampling positions and sample count. All 36 trials were eligible, yielding 839 pairs for across-participant evaluation. A sensitivity analysis excluded the participant contributing only four genuine central-rest task–baseline pairs per allocation and independently refitted the across-participant models in the remaining 24 participants (835 pairs).

The first 300 s of each participant’s baseline were divided into thirty nonoverlapping 10-s segments. These segments supplied the physiological comparison class. Each allocation used all available segments before reshuffling for reuse, independently of pupil values or classification outcomes. To characterize variability arising from baseline-segment assignment, 199 Monte Carlo donor allocations were evaluated. Reused segments remained grouped within participants.

Task and baseline segments underwent the same direct standardization or linear detrending followed by standardization. Gradients were computed on each complete transformed segment before central sample selection. A separate sensitivity recomputed transforms and gradients within the 2–8-s segment; full-rest and both central-rest specifications are reported in Supplementary Table 4. Twelve candidates crossed these transformations with eleven pupil-distribution features or 34 pupil-and-gradient features, and L1 logistic regression, L2 logistic regression or LDA. The same estimator settings were used for state and task–baseline classification (Supplementary Methods).

Nested across-participant evaluation held out one participant and used three participant-grouped inner folds. Nested within-participant evaluation used five outer and three inner GroupKFold splits, grouping baseline segments so that both pair members and all reuse of a segment remained together. Feature scaling was fit on training data only, and the model with the highest mean inner-fold AUC was selected within each allocation. The participant with four central-rest pairs could not support five outer donor-group folds; within-participant evaluation therefore included 24 participants (835 pairs), preserving the original fold requirements. Held-out predictions were used to calculate participant metrics.

Participant AUC and balanced accuracy were averaged across all 199 Monte Carlo baseline allocations before equally weighted group summaries. Conditional participant-bootstrap 95% CIs were calculated from 20,000 resamples of allocation-averaged participant metrics and therefore quantify participant uncertainty conditional on the fitted predictions. Allocation SDs and ranges separately quantified variability arising from baseline-segment assignment. Complete donor-allocation and model-selection records are provided in Supplementary Data 6.

### Exact-onset classification from preceding history

The onset analysis used a fixed LightGBM classifier in 25 participants. Target-replay onsets were identified on each full valid raster and then restricted to [2,8) s across all 36 trials. Each onset was matched to the same rest time in the nearest eligible other trial, using a cyclic search that considered the preceding trial before the following trial and required no target activity within ±250 ms of the matched sample. Matched control samples could be reused across onsets. The primary analysis comprised 12,063 pairs; Supplementary Methods describe the independently refitted 24-participant sensitivity.

Pupil histories were 50, 75, 100, 150, 200, 250, 300, 400 and 500 ms long, ending immediately before the candidate sample. Nineteen features summarized pupil level, variability, quantiles, endpoints, slope and first differences. Pupil was downsampled from 600 to 200 Hz and standardized by the pooled practice/rest mean and population SD within participant.

Sixty-four replay-history features summarized preceding onset counts, occupancy, recent onset and inter-onset lags, exponential lag summaries and consecutive-gap clusters. Eight context features encoded trial and rest position, early-trial and first-half indicators, and sine/cosine representations of trial and rest position. Exact feature definitions are given in Supplementary Methods; Supplementary Data 5 contains the machine-readable model and performance records.

LightGBM used 350 estimators, eleven leaves, learning rate 0.035, minimum child samples 30, column subsampling 0.9, L2 regularization 2 and L1 regularization 0.05. Four participant-held-out GroupKFold folds used training weights inversely proportional to participant sample counts, normalized to mean one. The designated 500-ms history window was specified before applying the model to the current cohort. Participant AUCs were equally weighted in group summaries, with 95% t intervals and one-sided t tests against 0.5. BH adjustment covered the nine pupil-only histories separately within each cohort. Paired feature-set comparisons quantified the contribution of each history type (Supplementary Fig. 7).

### Statistical analysis

The primary analyses examined behavioral learning, the association between pupil state and replay, slow pupil-phase organization of replay, and classification of replay-associated states and replay onset. Participant was the group inferential unit. Group means and effects used 95% t intervals unless a bootstrap or model-based interval is specified. Standardized paired effect size d_z_ was the mean participant difference divided by its sample SD. Tests were two-sided except for the explicitly stated one-sided above-chance classifier tests. Between-participant associations were evaluated using two-sided Pearson correlations. Within-participant regressions and occurrence models used participant-clustered uncertainty. Primary behavioral and between-participant association tests were reported without multiplicity adjustment. Multiplicity correction was applied within the explicitly defined sensitivity and control families described in the Supplementary Information. Eligibility criteria and resulting sample sizes are specified for each analysis.

## Data availability

The original behavioral, MEG and pupillometry recordings are publicly available through OpenNeuro (dataset ds006502, version 1.0.0) [18]. Processed data supporting the reported analyses and figures accompany the manuscript as Supplementary Data 1–6.

## Code availability

Custom analysis and figure-generation code accompanies the manuscript for review and will be made publicly available on GitHub upon publication. The published replay-estimation code is available at https://github.com/hcps-ninds/Replay [15].

## Author contributions

Debadatta Dash: Conceptualization, Methodology, Software, Formal analysis, Data curation, Visualization, Writing – original draft, Writing – review & editing. Leonardo Claudino, Hisato Sugata, Fumiaki Iwane: Validation, Writing – review & editing. Leonardo G. Cohen: Resources, Supervision, Funding acquisition, Project administration, Writing – review & editing. Ethan R. Buch: Resources, Data curation, Validation, Supervision, Writing – review & editing.

