## supplementary materials for "Constricted pupil states index neural replay during wakeful rest in humans"

*Supplementary Information*

### Supplementary Methods

#### Pupil preprocessing

Continuous pupil data were filtered before epoching using the Kret–Sjak-Shie Pupil Size Preprocessing Code v1.1 (Kret and Sjak-Shie, 2019). Exact zeros were replaced by the continuous-trace median, and the absolute trace minimum was added to the signal before filtering. The final smooth reconstruction supplied the filtered pupil trace. The preprocessed pupil trace was retained in arbitrary units (a.u.); subsequent analyses used analysis-specific within-participant normalization or baseline referencing.

Filtering rejected short isolated segments, excessive dilation speed, gap-edge artifacts and deviations from a smooth reconstruction. Islands shorter than 50 ms and separated by more than 40 ms were rejected. Dilation speed used the maximum absolute slope to neighboring valid samples, excluding gaps longer than 200 ms; the threshold was median speed + 16 median absolute deviations (MADs). Gaps longer than 75 ms and shorter than 2,000 ms received 50-ms edge padding. Up to four passes rejected absolute deviations from the reconstruction above their median + 16 MADs. MAD denotes the unscaled median absolute deviation from the median (Leys et al., 2013).

Valid samples were linearly interpolated to 100 Hz with nearest-neighbor edge extrapolation, smoothed with a first-order 16-Hz Butterworth filter applied forward and backward, and interpolated back to the original sample times. Bidirectional smoothing can spread post-onset information into preceding measurements; onset histories therefore support temporal discrimination rather than online forecasting. Analysis-specific normalization is defined in the main Methods.

#### Temporal controls

The fixed-label temporal control rotated the 12 central residual-pupil bins within each valid early trial while preserving trial means and replay-bin counts. The 20,000 draws included identity rotations and used shared offsets for target and control. For studentized contrasts, two-sided empirical p values were calculated from the conditional reference distributions using absolute deviations from zero and a plus-one correction. These conditional reference distributions assess additional temporal alignment given trial structure; they do not establish cyclic stationarity. The primary $N = 25$ result and eligible $N = 24$ cohort sensitivity are reported in Supplementary Fig. 3C,D. The sensitivity omitted the same sparsely supported participant as the state-effect sensitivity and computed the studentized statistic using the same 20,000 stored participant-level null draws.

Phase controls matched absolute rest time in two other early trials without same-raster activity within ±50 ms. Matching and participant averaging followed the main Methods. Full- and central-rest phase comparisons were evaluated (Supplementary Fig. 4). Benjamini–Hochberg (BH) correction covered four phase comparisons.

**Classification metrics and validation**

State classification used neural replay labels to select pupil samples, with one feature pair per eligible trial. Area under the receiver operating characteristic curve (ROC AUC) ranks class scores within a held-out participant; paired accuracy is the fraction of matched pairs in which the replay-associated vector scores higher; balanced accuracy averages class-specific accuracies at threshold 0.5. Across-participant evaluation holds out all observations from a participant; within-participant evaluation keeps each trial pair together. These metrics characterize different aspects of discrimination under nested and fixed-model evaluation (Supplementary Fig. 8).

Nested evaluation selected models using only outer-training data; fixed-model matching-scale comparisons used the same extended-feature linear discriminant analysis (LDA) at 0.5, 1, 1.5 and 2 s. Group intervals are participant 95% t intervals. Mean ROC curves were calculated by equally weighting participant-level ROC curves. Model-selection results and held-out scores are provided in Supplementary Data 5.

The compact state-classification representation contained 11 features of selected pupil samples: mean, sample SD, skewness, excess kurtosis, root-mean-square value, minimum, maximum, range, median, and 25th and 75th percentiles. Skewness was the mean cubed deviation divided by population SD cubed plus $10^{-12}$; excess kurtosis was the mean fourth-power deviation divided by population variance squared plus $10^{-12}$, minus three. The extended representation added 10-bin histogram entropy ($-\Sigma\ p \log p$ over occupied bins, with natural logarithms), the 5th, 10th, 90th and 95th percentiles, and the unscaled median absolute deviation, giving 17 features. Applying these same 17 summaries to the pupil gradient yielded 34 features in total. Percentiles used linear interpolation.

Each complete 200-Hz rest trace was either directly standardized or first linearly detrended by least squares over equally spaced positions from −1 to 1, then standardized by its sample SD plus $10^{-12}$. Gradients were computed on the complete transformed trace using central differences internally and one-sided differences at the endpoints, with unit sample spacing, before replay-associated or matched samples were selected within [2,8) s. Absolute-time matching strata were intersected with that interval, retaining partial boundary strata for the fixed 1.5-s sensitivity. The deterministic sample-selection seed was 42 + 100000s + 1000t + a, where s and t are zero-based participant and trial indices and a is the stratum start sample. All selected samples from eligible strata within a trial were pooled to form each feature vector.

State and task–baseline classifiers used training-fold StandardScaler transformations (mean centering and population-SD scaling). Logistic regression used L1 or L2 regularization, C = 1, the liblinear solver, balanced class weights, an intercept, tolerance $10^{-4}$, at most 2,000 iterations and random state 42. LDA used the singular-value-decomposition solver, tolerance $10^{-4}$, empirical class priors and no shrinkage. Candidate models were evaluated in the declared grid order; ties in mean inner-fold AUC were resolved by selecting the first candidate in that order. GroupKFold splits were deterministic and unshuffled. The task–baseline grid used the same two transformations, two feature families and three estimators; baseline gradients were computed before selecting the same relative sample positions as the paired task trace.

#### Task-replay pupil versus pre-task baseline pupil

The central-rest task–baseline comparison included N = 25 and 839 trial pairs for across-participant evaluation; the separately refitted N = 24 sensitivity included 835 pairs. The omitted participant contributed four genuine pairs per allocation and could not support the unchanged five-fold within-participant evaluation, which therefore used N = 24. Participant metrics were averaged over all 199 Monte Carlo baseline allocations before group summaries. Supplementary Table 1 reports performance and uncertainty; Supplementary Fig. 6 shows across-participant distributions and allocation variability.

Conditional participant-bootstrap 95% intervals used 20,000 resamples of allocation-averaged participant metrics for both validation designs. They quantify participant uncertainty conditional on fitted predictions, without full model-retraining or overlapping-training-set uncertainty. Allocation SDs and ranges separately quantify donor-assignment variability across the 199 Monte Carlo donor allocations of the same recordings.

For each cohort and validation design, bootstrap samples drew participants with replacement to the original cohort size, and the sampled allocation-averaged metrics were then averaged with equal participant weight. The 2.5th and 97.5th percentiles of 20,000 bootstrap means defined the conditional interval, using NumPy default_rng with fixed seeds 20264300 for across-participant and 20260925 for within-participant estimates. The same sampled participant indices were used for AUC and balanced accuracy. Allocation SDs used the sample SD across the 199 allocation-level group means; allocation ranges were their minimum and maximum.

Baseline segments served as the physiological comparison class, with task and baseline sampling positions and counts matched within each pair. Reused donor segments remained grouped during validation. Because the trained sequence had not yet been introduced or practiced, pre-task rest served as a physiological reference in which experience-dependent replay of the trained sequence was not expected. Because pre-task rest and inter-practice rest occurred in different behavioral contexts, this comparison provides a physiological reference rather than a replay-specific control. Supplementary Data 6 contains the allocation, donor and model-selection records.

#### Preceding-history onset classification

The fixed LightGBM model was evaluated with participant-held-out folds in N = 25. One participant supplied four onset/control pairs; the N = 24 sensitivity excluded that participant from both training and evaluation and refitted the model. Supplementary Table 2 reports all nine pupil-history AUCs, participant 95% t intervals, one-sided *p*-values and within-cohort BH q-values. The 500-ms pupil result did not survive the nine-window correction in either cohort. Supplementary Fig. 7 compares pupil, replay and context histories and the descriptive change in AUC when pupil features were added. Supplementary Data 5 contains the machine-readable model specifications and held-out performance records.

For each history length h, the 19 pupil features were the mean, population SD, minimum, maximum, range, 10th, 25th, 50th, 75th and 90th percentiles, first value, last value, last-minus-first difference, least-squares slope against time in seconds, and the mean, population SD, minimum, maximum and mean absolute value of successive differences. The window contained max(3, round(200h)) samples ending immediately before the candidate sample; any shorter vector was left-padded with its first value. Quantiles used linear interpolation. Features were calculated from participant-standardized pupil traces without additional within-window standardization. Nonfinite standardized rest values were linearly interpolated in both directions, and any remaining missing values were filled with zero.

The 64 replay-history features comprised 34 onset-count and occupancy measures, six recent-onset lags, five recent inter-onset intervals, ten exponential lag sums, seven consecutive-gap cluster sizes, and the total onset count and count/h. Count and occupancy windows were 25, 50, 75, 100, 125, 150, 175, 200, 225, 250, 300, 400, 500, 750, 1,000, 1,500 and 2,000 ms, each capped at h and converted to at least one 200-Hz sample. Occupancy was the fraction of active raster samples. Onsets were restricted to the preceding h seconds; lags and inter-onset intervals were ordered from most recent, with unavailable values set to h + 0.005 s. Exponential sums were Σ exp(−lag/τ), with τ = 25, 50, 75, 100, 150, 200, 300, 500, 750 and 1,000 ms; an empty history gave zero. Consecutive-gap cluster size counted the latest onset and successive preceding onsets separated by no more than 100, 150, 200, 250, 300, 400 or 500 ms, giving zero if no onset was present.

The eight context features were trial/36, rest time/10, indicators for trials ≤11 and ≤18, sin(2π × rest time/10), cos(2π × rest time/10), sin(2π × trial/36) and cos(2π × trial/36), using one-based trial number and rest time in seconds. LightGBM used gradient-boosted decision trees for binary classification with the fixed settings reported in the main Methods, unrestricted tree depth, random state 42 and one computational thread. No additional feature scaling or model selection was applied. Feature arrays were stored as 32-bit floats. For training participant s, each observation received weight proportional to $1/n_s$, where $n_s$ was that participant's training-observation count; weights were normalized to mean one.

**Behavioral and gaze controls**

Early replay–behavior controls used the same N = 24 participants and frozen trials 1–11 measures as main Fig. 3A–C. Pearson correlations compared target and control replay rates with early offline gain and early online change; the original two-test target/control BH family was applied separately to each behavioral outcome. Early offline gain summed the ten transitions linking trials 1–11, excluding 11→12. Replay regularity was lower variability in within-trial inter-event intervals, quantified by IEI CV with the original event-support criterion. Spearman correlations and leave-one-participant-out Pearson correlations were descriptive robustness checks; their p-values are nominal and do not form new confirmatory families. Correlation CIs used the original 2,000-resample participant-bootstrap percentile procedure (seed 1209).

Target-rate Spearman correlations were ρ = 0.533 with early offline gain (p = 0.007) and −0.508 with early online change (p = 0.011); IEI CV versus early offline gain gave ρ = −0.519 (p = 0.009). Control-rate Pearson correlations were 0.121 with early offline gain (p = 0.572, q = 0.572) and −0.069 with early online change (p = 0.749, q = 0.749). These separate tests do not establish a target–control difference. Across leave-one-participant-out (N = 23 each), target-rate/offline r ranged 0.414–0.574 (p = 0.004–0.049), target-rate/online r ranged −0.574 to −0.380 (p = 0.004–0.073), and IEI/offline r ranged −0.589 to −0.394 (p = 0.003–0.063). Directions were retained, while nominal significance varied for online and IEI associations. Full numerical records, including rank CIs and control sensitivities, accompany Supplementary Data 1. Supplementary Data 1 includes the corresponding participant-influence records.

Low-motion pupil analyses restricted early central-rest bins by gaze speed alone or jointly by speed, dispersion and excursion; the 16-test BH family covered two rasters, two restriction rules and four cutoffs (Supplementary Fig. 5). Within each participant, bins were retained at or below the 50th, 60th, 70th or 80th percentile of gaze speed alone, or of all three motion measures jointly. Bins required at least 95% finite horizontal and vertical gaze samples. Speed was the median successive-sample Euclidean displacement multiplied by 600 Hz; dispersion was the Euclidean norm of horizontal and vertical population SDs; excursion was the Euclidean norm of their ranges. Participants required at least five replay-positive and five replay-negative retained bins. Numerical results are provided in Supplementary Data 3.

#### Early versus post-early pupil-state sensitivity

The 500-ms state contrast was calculated separately for trials 1–11 and 12–36. Both periods retained the original participant-specific exact-time profile estimated across all 36 trials and the sample SD across all rest bins. Invalid sentinel trials were excluded from event and comparison-bin selection. Replay-positive minus replay-negative means were calculated within each participant and period; participants required both classes in both periods. Two-sided paired t tests compared early with post-early effects. The central 2–8-s window was the primary period comparison, and full 0–10-s rest was a secondary sensitivity; BH q values additionally summarize the two window tests. Early learning contributed 260 valid trials (7–11 per participant), 2,166 replay-positive and 954 replay-negative central bins; post-early learning contributed 625 valid trials (25 each), 5,014 positive and 2,486 negative bins. Each participant was weighted equally.

All 25 participants contributed both periods. The central-rest mean effect was −0.129 SD early and 0.004 SD post-early; the paired difference was −0.133 SD (95% CI [−0.280, 0.014], $t(24) = -1.874$, $d_z = -0.375$, $p = 0.073$, two-window BH $q = 0.146$). For full rest, the means were −0.081 and 0.025 SD, respectively (paired difference −0.106 SD, 95% CI [−0.266, 0.055], $t(24) = -1.362$, $d_z = -0.272$, $p = 0.186$, $q = 0.186$). The direct period comparisons therefore do not support an early-specific effect; the significant early estimate and nonsignificant post-early estimate alone do not establish a period difference.

#### Common-sample early-learning associations and sensitivity

Figure 3A–C uses N = 24 sample with estimable early IEI CV. Early offline gain sums the ten transitions linking trials 1–11; early online change sums within-trial changes over trials 1–11. The physiological and within-participant behavioral analyses retain the original N = 25 cohort.

#### Exploratory sample-support sensitivity

The primary early-rest pupil-state analysis weighted participants equally. The excluded participant in the previous analysis contributed one target-replay-positive 500-ms bin during early rest (one detection over 110 s of valid rest exposure), limiting the support for this participant's replay-associated pupil mean. Omitting this participant as an exploratory sample-support check yielded a mean replay-minus-no-replay pupil difference of −0.081 SD (95% t CI [−0.151, −0.011], $p = 0.025$; N = 24), compared with −0.129 SD (95% CI [−0.249, −0.009], $p = 0.036$; N = 25) in the primary analysis. Thus, the direction was retained while the estimated magnitude decreased. This participant had no invalid replay trials or recorded behavioral malfunction flags and thus remained in the primary cohort; this check introduces no new minimum-event exclusion rule.

### Supplementary Data

Supplementary Data 1–6 provide the numerical source data and analysis outputs corresponding to the main and supplementary figures and tables.

### Supplementary Table 1. Nested task–baseline performance and allocation variability

Conditional participant-bootstrap 95% intervals condition on fitted predictions after averaging allocations. Allocation SD and range describe repeated donor assignments, not independent experiments. Balanced accuracy uses a probability threshold of 0.5. All 199 Monte Carlo allocations were included for each validation design and cohort.

#### ROC AUC

| Validation | N | Mean | Conditional participant-bootstrap 95% interval | Allocation SD | Allocation range |
|---|---|---|---|---|---|
| Across | 25 | 0.664 | 0.615–0.715 | 0.0137 | 0.631–0.702 |
| Across | 24 | 0.665 | 0.614–0.719 | 0.0096 | 0.638–0.688 |
| Within | 24 | 0.706 | 0.661–0.752 | 0.0144 | 0.665–0.747 |

#### Balanced accuracy

| Validation | N | Mean | Conditional participant-bootstrap 95% interval | Allocation SD | Allocation range |
|---|---|---|---|---|---|
| Across | 25 | 0.628 | 0.593–0.665 | 0.0107 | 0.598–0.653 |
| Across | 24 | 0.631 | 0.593–0.671 | 0.0083 | 0.609–0.651 |
| Within | 24 | 0.668 | 0.630–0.706 | 0.0133 | 0.626–0.703 |

Complete donor-allocation records are provided in Supplementary Data 6.

### Supplementary Table 2. Pupil-only onset inference across all preceding histories

Participant is the unit of inference. Intervals are participant 95% t intervals; *p* is one-sided against AUC 0.5; q is BH-adjusted across the nine pupil-only windows separately in each cohort. The N = 24 values come from a separate refit, not deletion of one participant's score from the primary model.

#### Primary N = 25

| History (ms) | AUC | 95% interval | Raw *p* | BH q (nine) |
|---|---|---|---|---|
| 50 | 0.5080 | 0.4926–0.5234 | 0.1468 | 0.3302 |
| 75 | 0.5155 | 0.5035–0.5275 | 0.0067 | 0.0602 |
| 100 | 0.4982 | 0.4736–0.5228 | 0.5581 | 0.7190 |
| 150 | 0.4985 | 0.4772–0.5197 | 0.5592 | 0.7190 |
| 200 | 0.5093 | 0.4975–0.5212 | 0.0584 | 0.1752 |
| 250 | 0.4925 | 0.4670–0.5180 | 0.7246 | 0.7246 |
| 300 | 0.4965 | 0.4766–0.5163 | 0.6412 | 0.7214 |
| 400 | 0.5031 | 0.4762–0.5300 | 0.4080 | 0.7190 |
| 500 | 0.5211 | 0.5025–0.5397 | 0.0138 | 0.0622 |

#### Refitted N = 24 sensitivity

| History (ms) | AUC | 95% interval | Raw *p* | BH q (nine) |
|---|---|---|---|---|
| 50 | 0.5099 | 0.4952–0.5245 | 0.0882 | 0.1322 |
| 75 | 0.5078 | 0.4970–0.5186 | 0.0748 | 0.1322 |
| 100 | 0.5099 | 0.4982–0.5216 | 0.0462 | 0.1042 |
| 150 | 0.5072 | 0.4948–0.5195 | 0.1210 | 0.1556 |
| 200 | 0.5125 | 0.4978–0.5273 | 0.0463 | 0.1042 |
| 250 | 0.5038 | 0.4856–0.5220 | 0.3335 | 0.3673 |
| 300 | 0.5028 | 0.4858–0.5198 | 0.3673 | 0.3673 |
| 400 | 0.5190 | 0.5016–0.5364 | 0.0169 | 0.0760 |
| 500 | 0.5202 | 0.5033–0.5372 | 0.0107 | 0.0760 |

### Supplementary Table 3. Participant flow

| Stage | Participants | Definition |
|---|---|---|
| Parent-study enrollment | 33 | Healthy adults enrolled in the parent study |
| Technical MEG exclusions | 2 | MEG-system problems during acquisition |
| Participants in released dataset | 31 | Available after two technical MEG exclusions from 33 enrolled participants |
| Excluded from current analysis | 6 | Replay, pupil or alignment exclusions described below |
| Primary cohort retained | 25 | Primary multimodal pupil–replay cohort |

Thirty-three participants were enrolled in the parent study; two were excluded because of technical problems during MEG acquisition, leaving 31 in the released dataset. Six were excluded from the current analysis: one had entirely nonfinite replay rasters; two had no target detections on valid trials; one had unusable pupil recordings and absent target/control replay detections; one had unusable pupil recordings; and one had flat or absent pupil/trigger channels that prevented reliable multimodal alignment. The primary cohort comprised 25 participants. Individual trials with invalid replay data were removed without excluding otherwise usable participants.

**Supplementary Table 4. Full-rest and central-rest classification**

Mean participant ROC AUCs. Central sampling retains the original whole-10-s transforms and gradients, then selects samples within [2,8) s. Central transforms recompute these quantities within [2,8) s and are a separate sensitivity. These are descriptive specification comparisons, not tests of a window interaction.

| Task / validation | N | Full rest 0–10 s | Central sampling 2–8 s | Central transforms 2–8 s |
|---|---|---|---|---|
| Nested state, across | 24 | 0.846 | 0.857 | 0.876 |
| Nested state, within | 24 | 0.789 | 0.816 | 0.818 |
| Fixed state, 0.5 s, across | 24 | 0.786 | 0.806 | 0.812 |
| Fixed state, 0.5 s, within | 24 | 0.701 | 0.739 | 0.727 |
| Fixed state, 1 s, across | 24 | 0.823 | 0.841 | 0.850 |
| Fixed state, 1 s, within | 24 | 0.745 | 0.749 | 0.750 |
| Fixed state, 1.5 s, across | 24 | 0.834 | 0.844 | 0.846 |
| Fixed state, 1.5 s, within | 24 | 0.762 | 0.750 | 0.766 |
| Fixed state, 2 s, across | 24 | 0.846 | 0.852 | 0.868 |
| Fixed state, 2 s, within | 24 | 0.765 | 0.773 | 0.785 |
| Baseline, across | 25 | 0.680 | 0.664 | 0.537 |
| Baseline, across | 24 | 0.694 | 0.665 | 0.536 |
| Baseline, within | 24 | 0.747 | 0.706 | 0.626 |
| Baseline, within | 25 | 0.739 | Not estimable[1] | Not estimable[1] |

State classification used 840 full-rest and 835 central-rest pairs (N = 24). Baseline classification used 846/840 full-rest pairs for N = 25/24 and 839/835 central-rest pairs. [1]The participant with four central-rest pairs cannot support the unchanged five outer folds; within-participant central estimates therefore use N = 24. All baseline specifications average 199 allocations. Exact-onset candidate times were already restricted to [2,8) s and that analysis is unchanged. Full-rest phase analyses are also unchanged.

Recomputing transforms within central rest reduced task–baseline discrimination (across N = 25 AUC 0.537, 95% interval 0.505–0.569; within N = 24 AUC 0.626, 0.587–0.668), showing sensitivity of this physiological-reference comparison to the transformation interval. Nested state discrimination remained strong under both central specifications. Machine-readable comparisons and held-out results are in Supplementary Data 5–6.

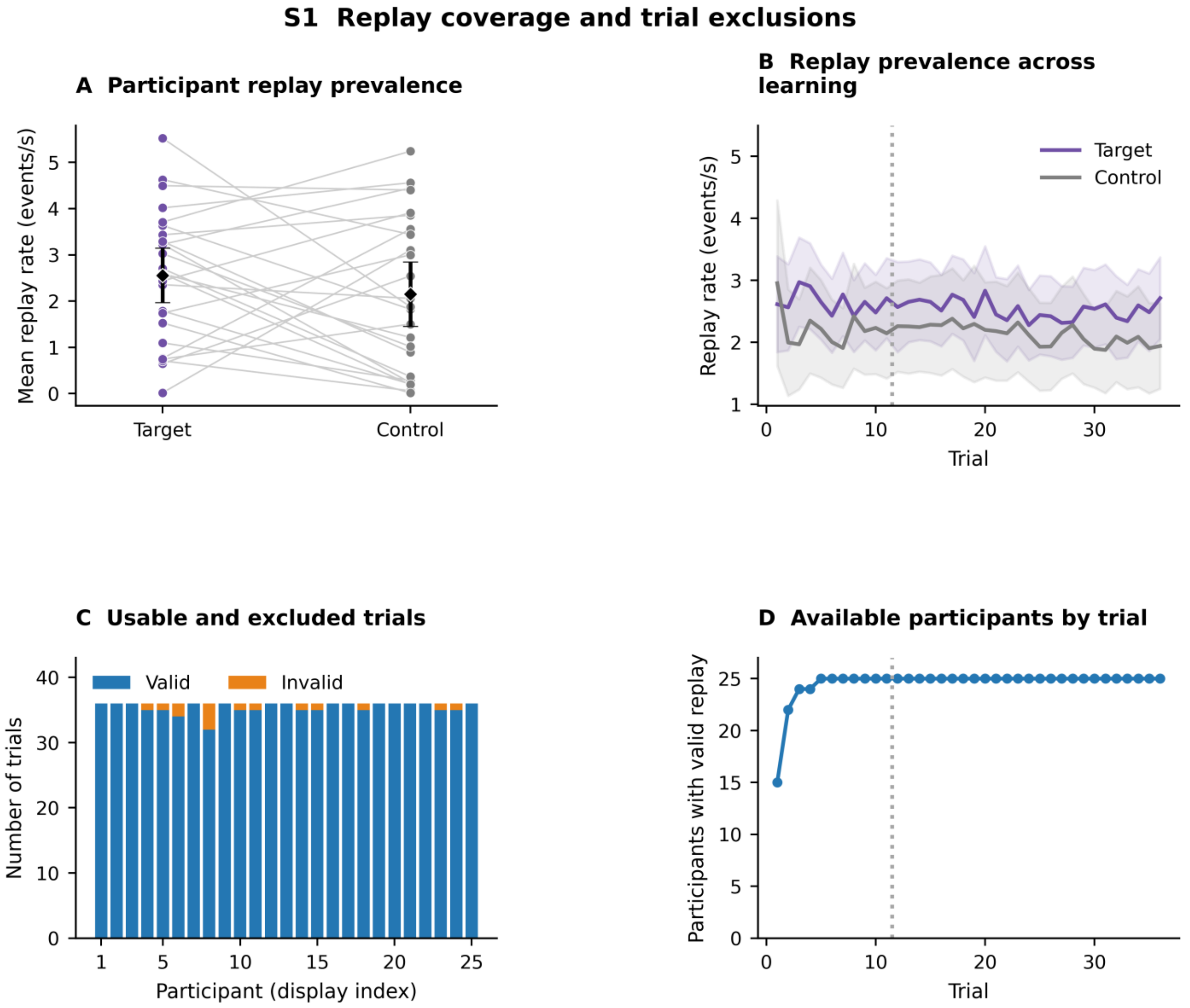


**Supplementary Fig. 1. Replay coverage and trial exclusions.** (A) Participant replay prevalence. Paired target and control rates show replay across the retained participants. (B) Replay prevalence across learning. Trial-wise rates show replay throughout training; the dotted boundary separates early trials 1–11 from post-early trials 12–36. (C) Usable and excluded trials. Valid and invalid trial counts identify the trial-level exclusions within otherwise usable participants. (D) Available participants by trial. Trial 1 included 15 participants and trial 2 included 22; the remaining 10 and 3 records, respectively, had invalid sentinel trials that contributed neither events nor exposure. These trial exclusions did not remove participants from the cohort. N = 25 overall; display indices identify participants consistently across panels. Trials with both rasters entirely active were invalid; an active first sample in an otherwise valid raster counted as an onset. Diamonds and bars in A denote means and 95% t confidence intervals (CIs); bands in B denote participant 95% t CIs. Numerical coverage tables are provided in Supplementary Data 3.

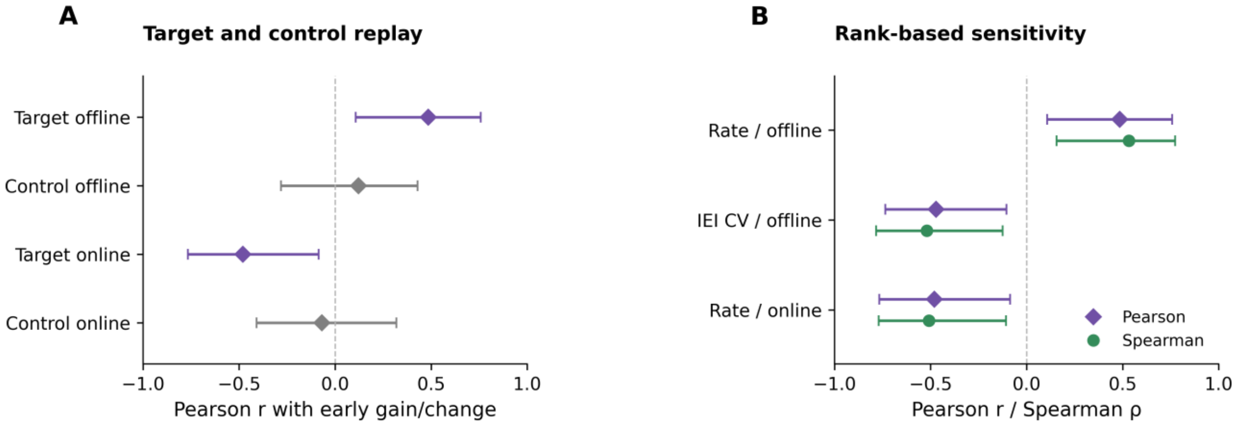


**Supplementary Fig. 2. Early replay–learning associations.** (A) Target and control early replay rates versus early offline gain and online change ($N = 24$). Target Pearson $r = 0.484$ ($p = 0.016$, $q = 0.033$) and $-0.481$ ($p = 0.017$, $q = 0.035$); control $r = 0.121$ ($p/q = 0.572$) and $-0.069$ ($p/q = 0.749$), respectively. BH correction covers the target/control pair separately for each outcome. (B) Pearson and Spearman estimates for the three main early-learning associations ($N = 24$). Lower IEI CV denotes less variable within-trial event spacing. Early offline gain uses the ten transitions linking trials 1–11; early online change covers trials 1–11. A/B show 2,000-resample participant-bootstrap percentile 95% CIs. Rank and influence checks are descriptive. Complete estimates, uncertainty and correction families, are supplied in Supplementary Data 1 and 3.

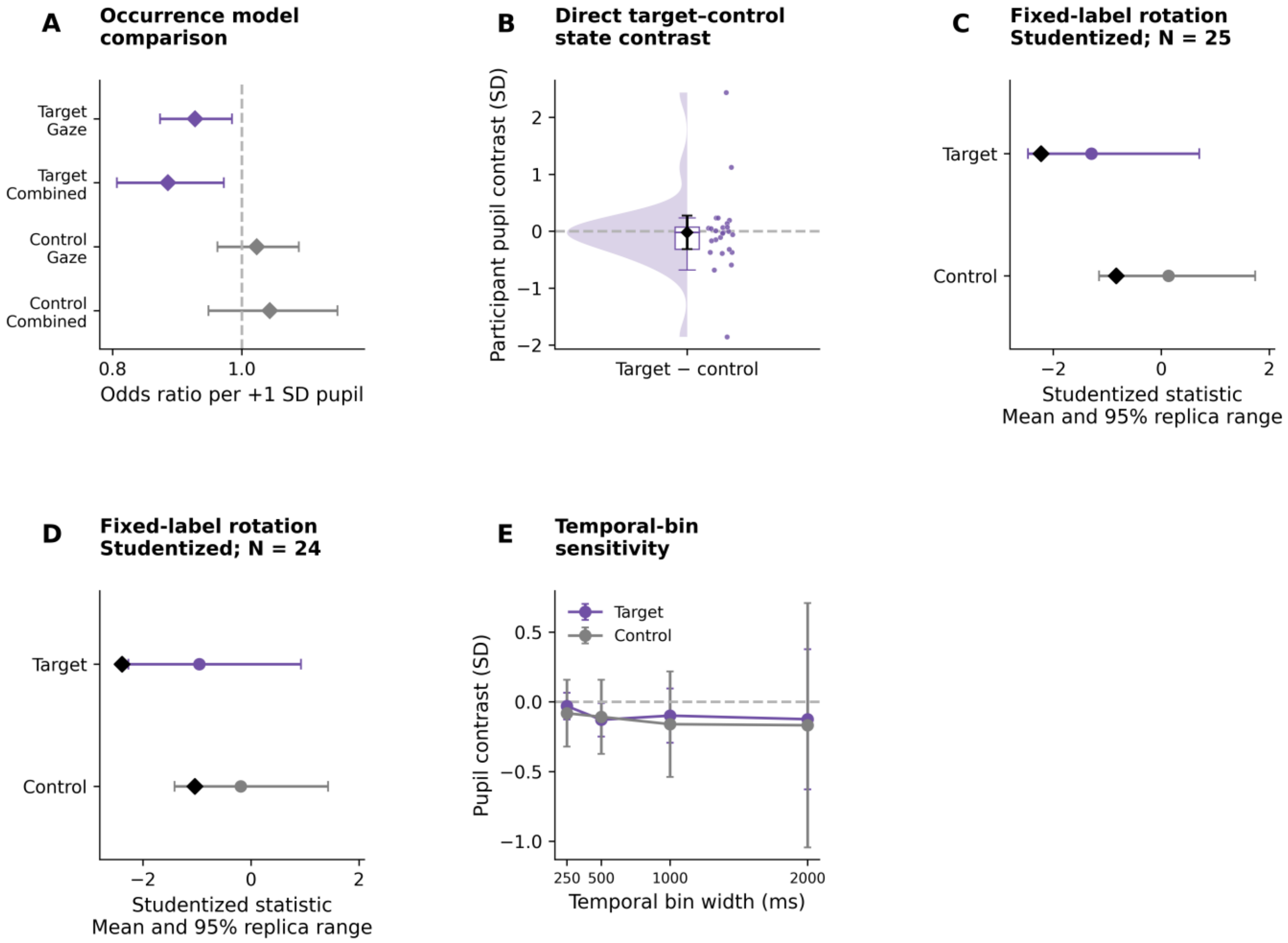


**Supplementary Fig. 3. Occurrence sensitivity and conditional temporal controls.** (A) Occurrence model comparison. Larger pupil was associated with lower target replay odds in gaze-adjusted and combined time-and-gaze models; the combined target test gave $p = 0.011$ and BH q = 0.021, whereas the control test gave $p = 0.382$ and q = 0.382. (B) Direct target–control state contrast. The participant target-minus-control contrast did not establish a sequence difference (paired $p = 0.883$). (C) Fixed-label rotation: studentized statistic, N = 25. The primary target result was inconclusive (empirical p = 0.073; control p = 0.344). (D) The corresponding exploratory N = 24 sample-support sensitivity gave target p = 0.015 and control p = 0.218, using the same 20,000 rotation draws after omitting the sparsely supported participant. (E) Temporal-bin sensitivity. Mean target contrasts at 250, 500, 1,000 and 2,000 ms were −0.031, −0.129, −0.099 and −0.125 SD, respectively; target N was 25, 25, 19 and 9 across these widths, with widening uncertainty at the coarser scales. The analysis in A used 3,120 bins, 260 trials and 25 participants, with participant-clustered 95% normal-Wald CIs and BH correction across two target/control tests. The combined coefficient includes within- and across-trial variation. Panels B and E use participant 95% t CIs. Panels C and D summarize reference means and central 95% ranges; black diamonds mark observed values. Rotations preserve trial means and replay-bin counts. In E, target 95% CIs were [−0.127, 0.065], [−0.249, −0.009], [−0.294, 0.096] and [−0.627, 0.377]; control N = 25, 25, 20 and 13. The control bin-level contrast was −0.108 SD ($p = 0.411$), and its fully adjusted OR was 1.044 (95% CI 0.948–1.148). Supplementary Data 2 contains the temporal-control results.

**S4 Phase specificity after matching rest time**

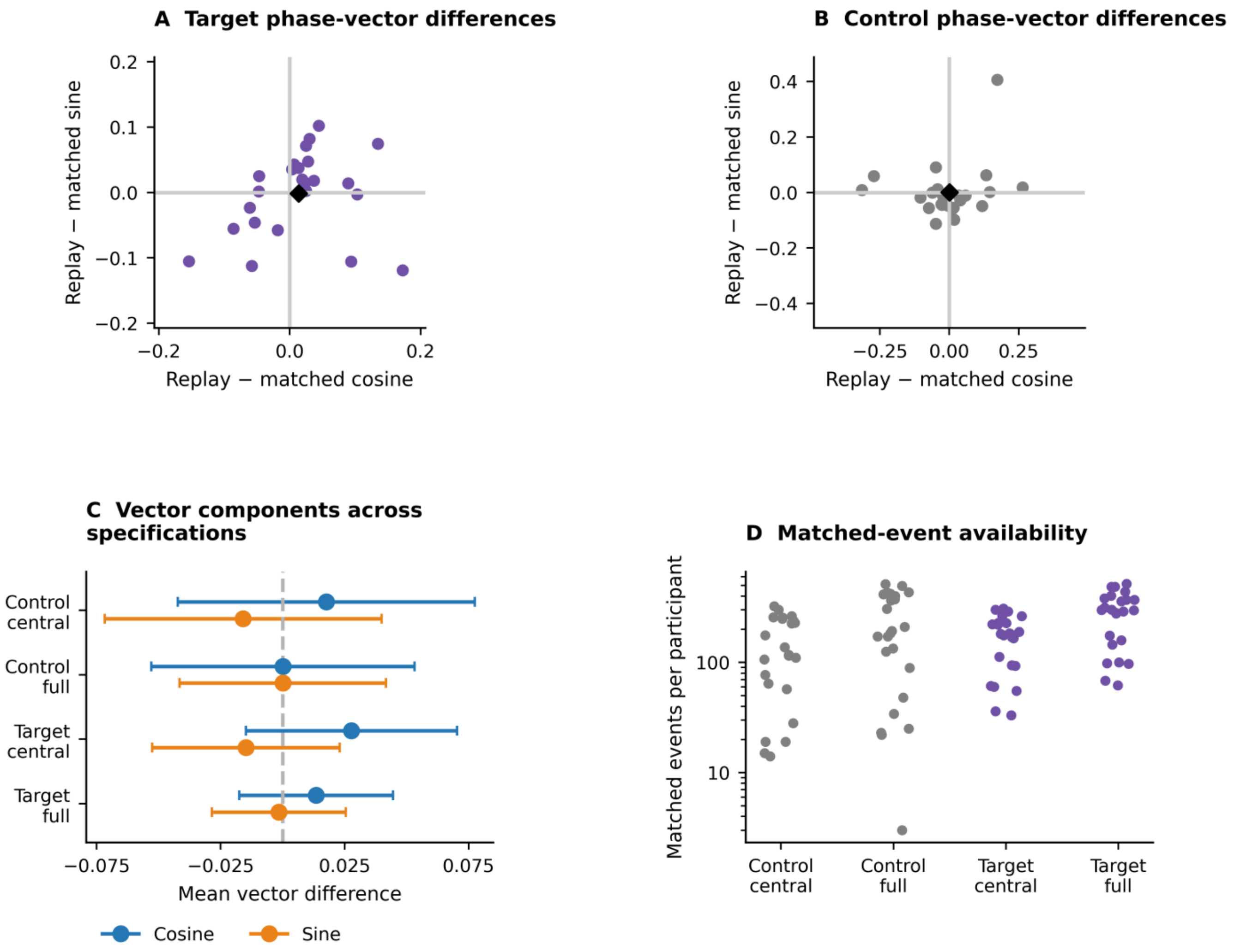


**Supplementary Fig. 4. Phase specificity after matching rest time.** (A) Target phase-vector differences. Participant-level target phase vectors relative to same-time controls in other early trials did not establish phase displacement beyond rest-time structure. (B) Control phase-vector differences. The corresponding control-replay comparison likewise did not establish displacement; black diamonds mark group means. (C) Vector components across specifications. Panel C shows cosine and sine differences for central and full rest for both replay types, with marginal participant 95% t CIs. (D) Matched-event availability. Matched-event counts on a logarithmic axis document the observations contributing to each comparison. Controls required two eligible other trials without same-raster replay within ±50 ms; participants required at least two matches. Two-dimensional Hotelling tests gave: control central rest, N = 23, $p$ = 0.621, q = 0.865; control full rest, N = 24, $p$ = 1.000, q = 1.000; target central rest, N = 24, $p$ = 0.307, q = 0.865; target full rest, N = 24, $p$ = 0.649, q = 0.865. BH correction covered these four tests. Control phase concentration before matching had Rayleigh $p$ = 0.026; matched null results do not establish equivalence. Slow phase uses continuous 0.1–1-Hz filtered pupil; full rest spans [0,10) s and central rest [2,8) s. Supplementary Data 3 provides individual vectors and matching counts.

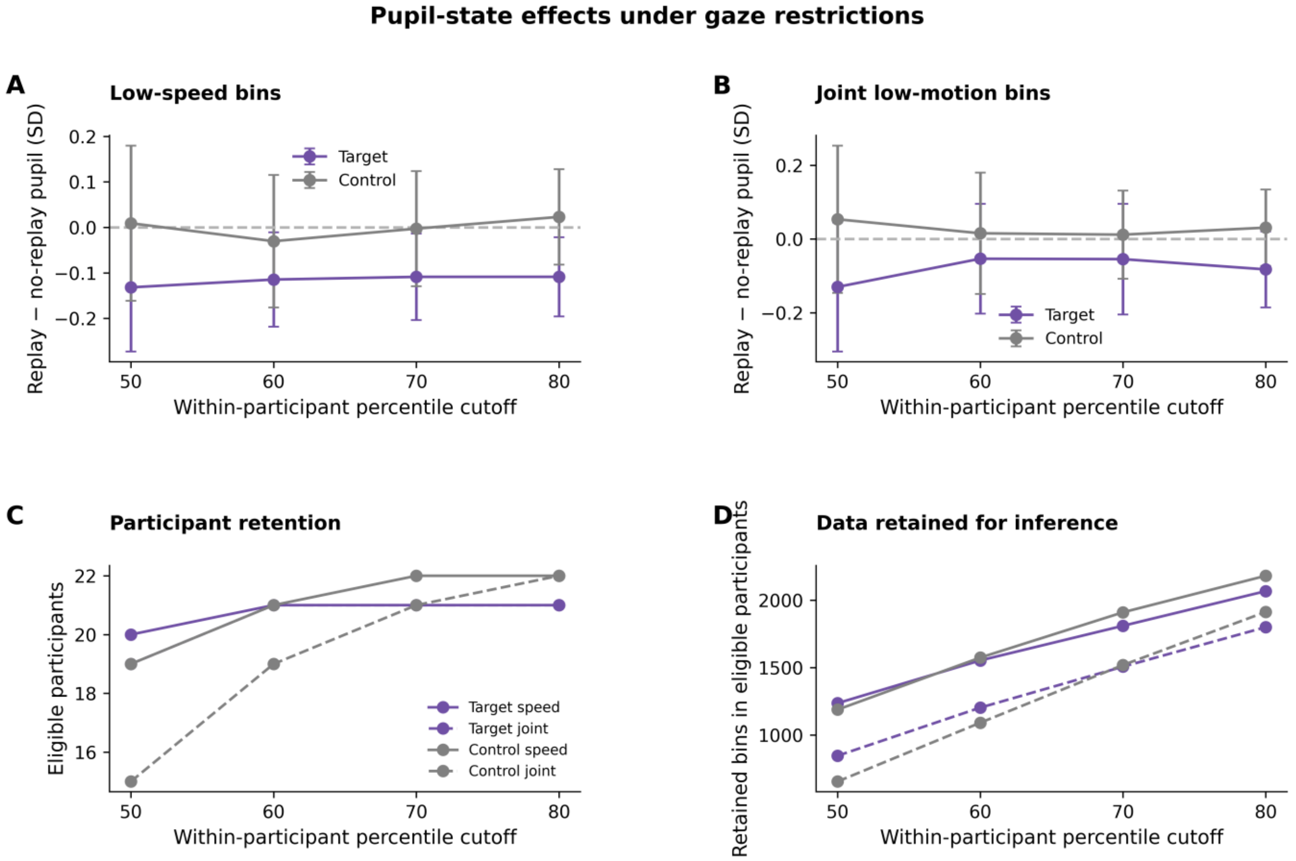


**Supplementary Fig. 5. Pupil state effects under gaze restrictions.** (A) Low-speed bins. Exact-time-residual pupil contrasts compare replay-positive and replay-negative bins after increasingly restrictive gaze-speed cutoffs. (B) Joint low-motion bins. Adding gaze-dispersion and excursion restrictions reduced the data available to estimate the pupil contrast. (C) Participant retention. Eligible counts fell under stricter restrictions, with N = 15–22 across comparisons. (D) Data retained for inference. Replay/no-replay bin counts quantify the reduction in observations under speed-only and joint restrictions. None of the 16 exploratory comparisons survived BH correction (all $q \geq 0.171$). All four within-participant cutoffs are displayed; panels A and B use participant 95% t CIs. Early central 500-ms bins require ≥95% finite paired gaze samples, and participants require ≥5 positive and ≥5 negative bins. Purple denotes target and gray control; solid lines denote speed-only and dashed lines joint restrictions. These restrictions identify reduced eye movement, not formally detected fixations or microsaccades; participant composition changes across restrictions. Supplementary Data 3 contains raw *p*-values and adjusted q-values.

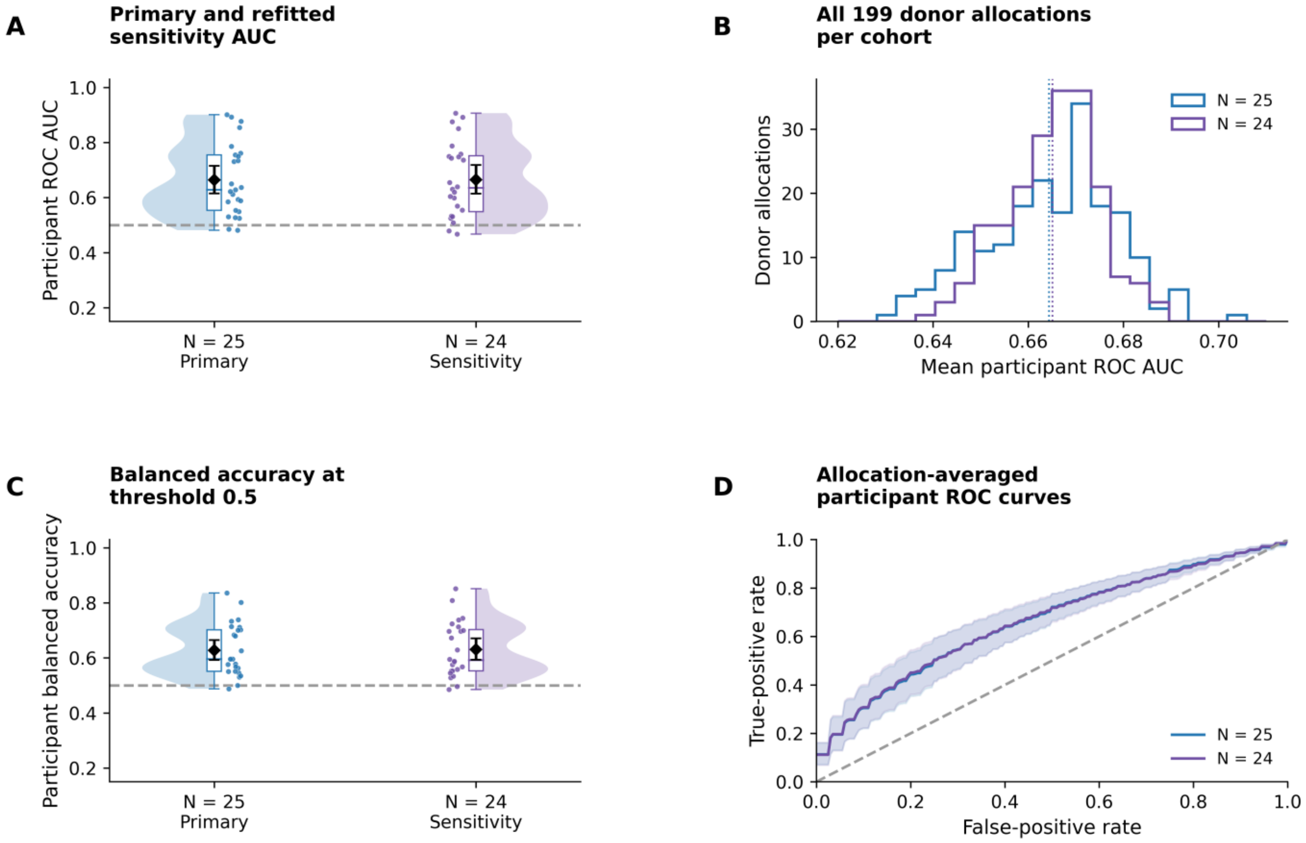


**Supplementary Fig. 6. Central-rest task-replay versus pre-task baseline classification.** (A) Primary and refitted sensitivity AUCs. Across-participant ROC AUC, averaged over 199 Monte Carlo baseline allocations, was 0.664 (95% interval 0.615–0.715; N = 25) and 0.665 (0.614–0.719; independently refitted N = 24). (B) Distributions of mean AUC across the 199 donor allocations quantify baseline-assignment variability, distinct from participant uncertainty. (C) Balanced accuracy at threshold 0.5 was 0.628 (0.593–0.665; N = 25) and 0.631 (0.593–0.671; N = 24). (D) Allocation-averaged participant ROC curves. Within-participant AUC was 0.706 (0.661–0.752; N = 24; Supplementary Table 1). Samples came from 2–8 s after whole-rest transformation. Dots represent participants; diamonds and bars represent means and conditional participant-bootstrap 95% intervals. Bands in D are pointwise conditional participant-bootstrap intervals; vertical lines in B mark allocation means. Supplementary Data 6 provides allocation-level and participant-level results.

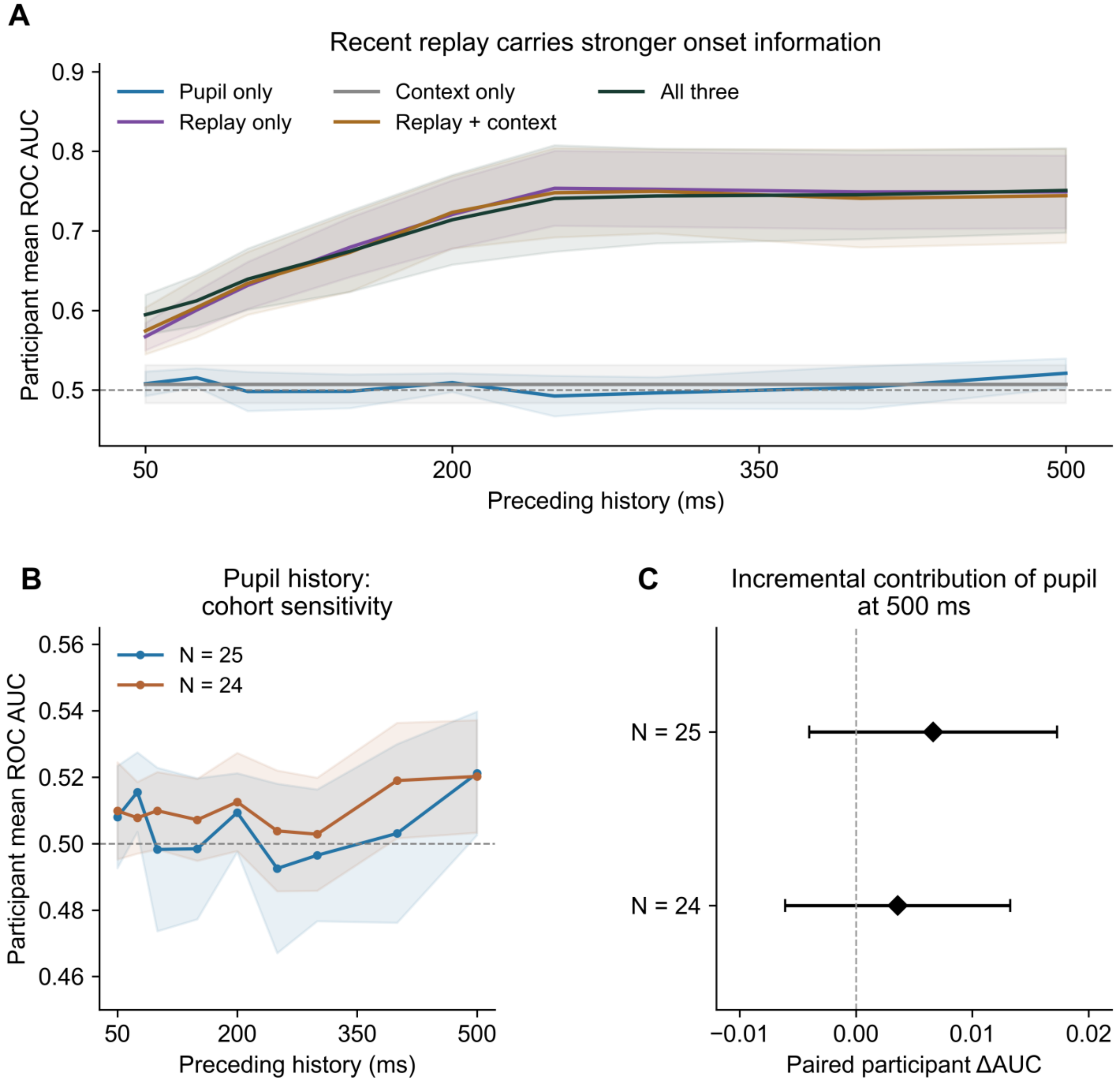


**Supplementary Fig. 7. Onset-history information and cohort sensitivity.** (A) Recent replay carries stronger onset information. Participant-held-out AUCs across nine histories compare pupil, replay, context and their combinations (N = 25); replay-only AUC reached 0.749 at 500 ms. (B) Pupil history: cohort sensitivity. Pupil-only discrimination was modest in both cohorts at 500 ms: AUC = 0.521 (95% CI 0.503–0.540), one-sided $p = 0.0138$ and nine-window BH q = 0.0622 for N = 25; the independently refitted N = 24 sensitivity gave 0.520 (0.503–0.537), $p = 0.0107$ and q = 0.0760. (C) Incremental contribution of pupil at 500 ms. Adding pupil to replay history and context yielded only a small descriptive paired ΔAUC: 0.00664 (95% CI [−0.00402, 0.01729]) for N = 25 and 0.00358 (95% CI [−0.00609, 0.01326]) for N = 24. Diamonds and bars represent means and participant 95% t intervals; bands in panels A and B are pointwise participant 95% t intervals. Supplementary Table 2 gives the complete nine-history inference, including multiplicity correction; Supplementary Data 5 contains source results.

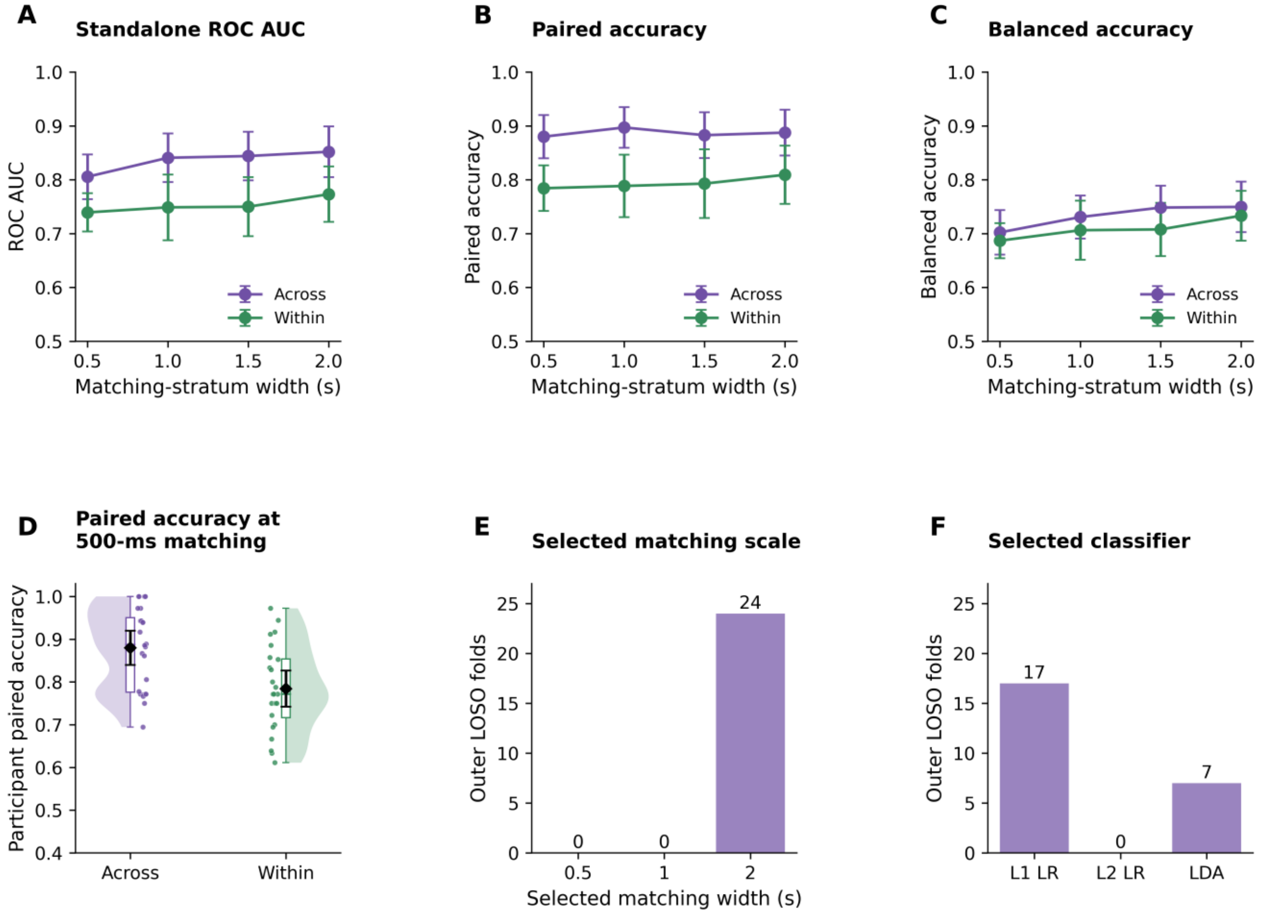


**Supplementary Fig. 8. Classification tasks and model selection.** (A) Standalone ROC AUC. The fixed model used linear detrending followed by trial standardization, the extended-feature representation and linear discriminant analysis (LDA) to characterize state discrimination across four matching widths. (B) Paired accuracy. Correct ranking of matched replay-associated and inactive observations summarizes paired discrimination across the same widths. (C) Balanced accuracy. Mean class-specific accuracy characterizes the same models at the classification threshold. (D) Paired accuracy at 500-ms matching. Mean paired accuracy was 0.880 across participants and 0.784 within participants. (E) Selected matching scale. All 24 outer nested folds selected 2-s matching. (F) Selected classifier. The central-rest nested search selected LDA in seven of 24 folds and L1 logistic regression in 17. All 24 folds selected the extended 34-feature representation; 20 selected direct trial standardization and four selected linear detrending followed by standardization. Panels A–D characterize fixed-model performance across matching scales, whereas E–F summarize model selections from the nested analysis. The 1.5-s width occurs only in the fixed-model sensitivity. Across-participant evaluation leaves one participant out (LOSO); within-participant evaluation groups whole trial pairs into five folds. N = 24; points represent participants, and diamonds and bars represent means and 95% t CIs. Supplementary Data 5 contains participant scores, held-out predictions and nested model-selection records.
